# Membrane partition and structural reorganization induced by anti-psychotics with distinct clinical profiles

**DOI:** 10.1101/2025.02.10.637357

**Authors:** Ana Gorse, Vesela Yordanova, Jessica Bodosa, Marion Mathelié-Guinlet, Astrid Walrant, Nada Taib-Mamaar, Axelle Grélard, Claire François-Martin, Rim Baccouch, Estelle Rascol, Gil-mar F. Salgado, Maria João Moreno, Margarida Bastos, Jeffery B. Klauda, Galya Staneva, Philippe Nuss, Isabel D. Alves

## Abstract

Antipsychotics (APs) are used in the treatment of severe mental disorders. Their mechanism of action involves interaction with multiple brain targets, notably the dopamine D2 receptors (D2R), where they compete with dopamine. Due to their lipophilic nature, APs also partition and accumulate in lipid membranes, particularly around the D2R and in synaptic vesicles. When intercalated into brain membranes, APs slowly accumulate and act as a reservoir, allowing their rapid release on demand to modulate neurotransmitter signaling. They also modify the physicochemical and mechanical properties of the lipid bilayer. These modifications can subsequently affect the conformational changes of embedded membrane proteins like the D2R. The present study investigated two major APs with different pharmacological and clinical profiles: chlorpromazine, which exerts its clinical activity mainly through a strong antagonistic action at the D2R, and clozapine, the weakest D2R antagonist of all APs. Surprisingly, although D2R antagonism is usually associated with AP potency, clozapine has repeatedly demonstrated clinical superior efficacy to all APs and is therefore recommended for treatment-resistant schizophrenia. The current work aims to extend the classical AP receptor mediated paradigmatic mode of action to their potential and unique membrane remodeling properties by thoroughly comparing their partitioning and impact on the physicochemical properties of the lipid membrane. Lipid model membranes mimicking synaptic vesicles have been investigated using a combination of several biophysical methods. The study aims to determine how the partitioning of the two APs modifies membrane order, phase transition, thickness, elasticity, phase separation, membrane integrity and charge. Differences have been demonstrated between these two compounds, which may further differ both over time as they accumulate as well as depending on their pre- or post-synaptic location.

## INTRODUCTION

Brain signaling is a vast scientific and conceptual field of investigation in neuroscience and psychiatry that covers approaches ranging from the molecular scale to the anthropological level, such as human consciousness. Among the various signaling modalities in the brain, those involving the lipid composition of neural cells are of considerable importance. This is due to the intrinsic role of brain lipids as actors of the message propagation, and because a considerable number of drugs, particularly those used to treat neuropsychiatric disorders, tend to accumulate in cell membranes and thus affect their biophysical properties. This phenomenon is observed in the case of antipsychotic drugs (APs), which specifically intercalate into the lipid membrane depending *e*.*g*. on their electrochemical and steric configuration, precise lipid composition and membrane organization.

Lipids represent a significant component of the brain, where they are known to act at various levels in terms of signaling. Brain lipids were primarily conceived as signal facilitators *e*.*g*. through the myelin membrane, a highly specialized lipid cell membrane indispensable for rapid nerve conduction. Recent data reinforced the idea that lipids play a more extensive role in the generation, transmission and hierarchization of brain information. This influence can occur through the action of single lipids extracted from the *in-situ* membrane, such as endocannabinoids, or by affecting the biophysical properties of the nerve cell membrane, and therefore depending on the nature, membrane location and organization of the lipids within the membrane.^1–3^ This is of particular importance at the synapse level, as *e*.*g*. lipidic secretory vesicles, which are in the presynaptic compartment, that are essential for transmitting chemical messages.

Neurons contain two primary classes of regulated pre-synaptic-located secretory lipid vesicles: large dense core vesicles (LDCVs), which store neuropeptide transmitters, and synaptic vesicles (SVs), which contain only neurotransmitters such as dopamine. Fusion of SVs with the pre-synaptic plasma membrane, releasing their neurotransmitter content, is balanced by endocytosis, which maintains the presynaptic membrane in a steady state. In particular, the extent to which this regulation is affected by the intercalation of psychotropic molecules, such as antipsychotics (APs), which are known to accumulate slowly in SV membranes, remains to be elucidated.^4^ The biophysical changes that APs induce in the lipid membranes of the SVs may be implicated in the control of the dopamine signaling, a role traditionally attributed to their binding to dopamine D2 receptors embedded in the postsynaptic membrane. The locoregional specialization of the lipid composition of the brain membrane also encompasses differences between pre- and post-synaptic compartments where the specific composition of lipid membranes at play in the signaling process are critical.^5,6^ This applies to the lipid composition of SV membranes, which differs from that of the postsynaptic ones. It has been shown that mammalian SVs are enriched in cholesterol (Chol) and unsaturated fatty acyl chains compared to the plasma membrane. As the smallest and thus most highly curved membrane vesicles in mammalian cells, SVs also contain critical phospholipids involved in curvature, such as phosphatidylethanolamine (PE) along with those involved in fusion, such as phosphatidylserine (PS).^7,8^ Typically, SVs contain 40 % of Chol, 17 % of phosphatidylcholine (PC), 20 % of PE, 6 % of PS and 3.6 % of sphingomyelin (SM).^9^

As previously stated, the biophysical changes that APs induce at the neuronal lipid membrane level may constitute an additional mechanism of action, in conjunction with their canonical receptor binding properties. The planar, often cationic, and amphiphilic nature of APs allows their intercalation into the lipid bilayer of cellular membranes, resulting in variations in the lateral and transverse organization of membrane lipids and the turnover of phospholipids, leading to changes in membrane composition, fluidity, thickness, curvature, and integrity.^10–12^

These AP-induced effects on the biophysical properties of neuronal membranes can affect signaling both by altering the conformational changes of embedded G-protein coupled receptors (GPCRs) through modifications of their membrane lipid environment, and by affecting the size and properties of synaptic vesicles, particularly their fusion with the presynaptic membrane.^13^ Moreover, it was shown that the association of the ligands with the membrane affects their apparent affinity for membrane proteins.^14^ Just as neurotransmitters can be both membrane-bound and free, APs can be incorporated into the lipid membranes in the vicinity of their GPCR, but also into the small aqueous compartment surrounding their receptor, where they can bind, dissociate and rebind.^6,15^ In addition, other properties of APs may play a role in modulating brain signaling, for example through gene regulation and methylation or by modifying neuronal microcircuitry and cortical oscillations.^16–18^

Moreover, it was demonstrated that different APs affect the lipid metabolism and may modulate the lipid composition by increasing the cellular content of acidic phospholipids, such as PS (as seen for chlorpromazine – CPZ), which can further promote its own interaction with the lipid membrane as seen by different approaches such as ssNMR, DSC, ITC.^19–21^ Indeed, in experiments of D2R reconstitution it has been shown that the presence of PS in the lipid mixture is important for the ligand interaction with the D2R.^22^ In addition, many APs are good antioxidants and decrease membrane lipid peroxidation.^23^ These effects might be of importance as lipid peroxidation has been shown to affect the affinity or number of binding sites in membranes for muscarinic, adrenergic and dopamine ligands.^24^

Recent research focuses on the presynaptic compartment that has been implicated in the physiopathology of schizophrenia.^25^ This is of interest because APs such as CPZ and CLOZ, two paradigmatic AP medications in the field of psychopharmacology, are known to slowly accumulate in the membranes of synaptic vesicles.^26^ Both compounds, as planar amphiphilic molecules, are known to induce changes in the biophysical properties of both pre- and post-synaptic membranes.^27^ CPZ, as most APs, exerts its clinical AP activity via a strong antagonistic activity at the post-synaptic dopamine D2 receptors, while CLOZ, one of the weakest D2 antagonists among APs is paradoxically the most clinically efficacious in treating schizophrenia, as evidenced by meta-analyses.^28,29^ The superiority of CLOZ has been primarily attributed to its distinctive receptor binding profile (and that of its active metabolite N-desmethyl CLOZ), which is characterized by a strong affinity for D4, M1 muscarinic, and glutamatergic N-methyl-D-aspartate receptors.^30^ An alternative hypothesis has been proposed based on the binding characteristics of CLOZ on the D2R or its interaction with adenosine A1 and D2 receptor heterodimers within the lipid membrane.^31^ CLOZ’s superiority may also be attributed to its comparatively limited chaper-one-like activity on the intracellular D2R pool. While several factors have been reported to explain CLOZ’s atypical AP behavior, including its remarkably different pharmacoperone activity, membrane remodeling properties of CLOZ and other APs have never been thoroughly compared.^32^

The present study, therefore, aims to extend the receptor-centered approach to AP action by investigating how CPZ and CLOZ affect the biophysics of lipid membranes. This encompasses revealing how they partition in the lipid membrane and how they modify membrane order, phase transition, thickness, elasticity, domain formation, integrity (permeability) and liposome size and charge. Understanding the nature of the membrane’s biophysical changes produced by different APs is important not only to investigate why APs with similar receptor binding profiles can produce different clinical responses, but also to provide insight into the unique nature of CLOZ. Here, we suggest that the superiority of CLOZ over CPZ may, at least in part, be related to its specific interaction with synaptic neuronal membranes and consequent membrane remodeling activity.

## EXPERIMENTAL SECTION

### Materials

Cholesterol (Chol), 1,2-dipalmitoyl-sn-glycero-3-phosphocholine (DPPC), 1-palmitoyl-2-oleoyl-glycero-3-phosphocholine (POPC), 1,2-dioleoyl-sn-glycero-3-phosphoethanolamine (DOPE), 1,2-dioleoyl-sn-glycero-3-phospho-L-serine sodium salt (DOPS) and 1,2-dipalmitoyl-2-(2H62) sn-glycero-3-phosphocholine (2H62-DPPC) were obtained from Avanti polar lipids (Alabaster, AL), either in powder form or dissolved in chloroform. Brain and egg yolk sphingomyelin (SM) and the fluorescent markers 6-dodecanoyl-N, N-dimethyl-2-napthylamine (Laurdan) and sulforhodamine B - acid form (SRB) were purchased from Sigma-Aldrich. The antipsychotics: clozapine (CLOZ) and chlorpromazine (CPZ) were both obtained from Sigma-Aldrich.

### Lipid model systems

A complex lipid model membrane mimicking the composition of the synaptic membrane was used (system 1). In addition, a simple model composed of POPC alone (system 2) and a third system composed of POPC and anionic lipids (system 3) were used to distinguish the potential role of electrostatic contribution (Table 1). The lipid model membranes were thus:

**Table 1.**
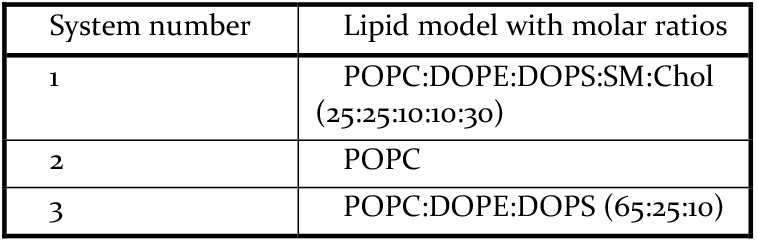
Lipid models used in the study.

- POPC:DOPE:DOPS:SM:Chol (25:25:10:10:30 molar ratios; system 1)
- POPC (system 2)
- POPC:DOPE:DOPS (65:25:10 molar ratios; system 3)

The three lipid systems were used in most approaches, exceptions depending on the limitations associated with each method, and/or to address specific questions.

### Preparation of Lipid Model Systems

Lipid films were prepared by dissolving appropriate amounts of lipids in chloroform or chloroform/ethanol (1/1 v/v) either in absence or presence of the antipsychotics (APs) chlorpromazine (CPZ) and clozapine (CLOZ). The organic solvent was evaporated under nitrogen flow at room temperature and then, in order to ensure the removal of all solvent traces, the samples were put in a desiccator under vacuum for at least 2 h. Then, lipid films were hydrated with the appropriate buffer, depending on the method, allowing the formation of multilamellar vesicles (MLVs). The MLVs were used to prepare both large unilamellar vesicles (LUVs) and small unilamellar vesicles (SUVs), for which experimental details are provided within the method applied.

### Marvin Sketch Simulations

Marvin Calculator Plugins were used for CPZ and CLOZ logP predictions and pKa microspecies distributions, Marvin 20.5.0, 2020, ChemAxon (http://www.che-maxon.com).

### Isothermal Titration Calorimetry

The heat flow resulting from titration of lipid vesicles (LUVs, 200 nm) into AP solution present in the cell was measured by isothermal titration calorimetry (ITC).^33^ The initial lipid concentrations for film preparation were 7 mM for the experiments with CPZ and 70 mM for CLOZ and the accurate lipid concentration following liposome extrusion was determined prior to the ITC measurement (detailed information of lipid preparation can be found in SI).^34^ CPZ concentration in the cell was 10 µM, and CLOZ concentration was 40 µM. Solutions of AP lipid suspensions were degassed for 10 min at 605 mmHg before the experiment. A VP-ITC instrument (Microcal/malvern) with a reaction cell volume of 1.4301 mL was used for the measurements and a total of 21 consecutive injections of 10 µL each were performed (the first was 4 µL and was discarder from the analysis), with 300 s interval between each injection. Measurements were performed at 25 °C. The heat of dilution was measured in separate experiments, by titrating the vesicles’ suspension into the buffer present in the cell and was subtracted for each AP/lipid titration data.

The thermograms were integrated using Origin 7.0 specifically modified by MicroCal for ITC measurements. The fitting and retrieval of partition thermodynamic parameters were then performed with Excel spreadsheet for analysis of ITC thermograms using the Solver add-in.^35–38^ The lipid molar volume value used was 0.795 dm^3^ mol^-1^ for POPC, DOPE and DOPS.^19^ The fraction of accessible lipid (γ) used was 0.5, considering lipid multilamellarity. In the presence of 10 % negative lipids with a charge of 1, a net membrane charge of - 0.1 was taken into account in the Excel spreadsheet data calculations.

### All-atom Molecular Dynamics Simulation

#### Parameterization of CPZ and CLOZ

CPZ and CLOZ were parameterized using the CGENFF module of CHARMM-GUI in their neutral, uncharged state.^39,40^ Free energy perturbation (FEP) was done in CHARMM to calculate the octanol-water partitioning free energy, ΔG_o/w_ of CPZ and CLOZ (the details are mentioned in the SI). The calculated ΔG_o/w_ with standard error (σ/N) and the logP value from the DrugBank portal are listed in Table 2. Since the calculated ΔG_o/w_ is close to the experimental value, we used the CGENFF parameters without any further changes.^41,42^

**Table 2.**
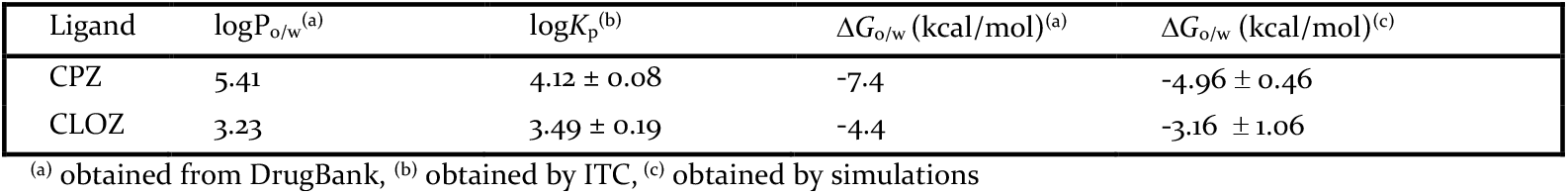
Free energy of ligand partitioning from water to octanol (ΔG_o/w_) obtained with simulations. LogP values obtained from DrugBank and logK_p_ values obtained with ITC (POPC vesicles, pH 7.4, 25 °C).

| Ligand | $\log P_{o/w}^{(a)}$ | $\log K_p^{(b)}$ | $\Delta G_{o/w}$ (kcal/mol)<sup>(a)</sup> | $\Delta G_{o/w}$ (kcal/mol)<sup>(c)</sup> |
| --- | --- | --- | --- | --- |
| CPZ | 5.41 | $4.12 \pm 0.08$ | -7.4 | $-4.96 \pm 0.46$ |
| CLOZ | 3.23 | $3.49 \pm 0.19$ | -4.4 | $-3.16 \pm 1.06$ |

(a) obtained from DrugBank, (b) obtained by ITC, (c) obtained by simulations

#### System Setup and Simulation Protocol

CHARMM-GUI’s *Membrane Builder* was used to set-up the three membrane-only systems.^43,44^ The membrane-only systems were equilibrated following the standard CHARMM-GUI protocol in namd 2.14.^45^ CHARMM-GUI’s Multi-component assembler module^46^ was used to prepare the equilibrated membranes with high (100:20 lipid:AP molar ratio) or low (100:9 lipid:AP molar ratio) concentration of APs equally spaced in water at a distance of 1 nm from the membrane (neutral form). Each system was therefore simulated in 5 different settings: a membrane-only (reference) simulation, two low AP concentration simulations and two high AP concentration simulations, all listed in Tables S2 and S3. The membrane-AP simulations were equilibrated with harmonic restraints (k=1000 kJ/mol) on the ligands which were later removed in the production runs done in GROMACS 2021.4.^47,48^ All simulations used the CHARMM36 forcefield for lipids and TIP3 water.^49–51^ All systems were first energy minimized and equilibrated in NVT and then the final production runs were done in NPT at 296.15 K. The production runs were done using the Nosé-hoover thermostat and Parinello-Rahman barostat with semi-isotropic coupling for the membrane. The LINCS algorithm was used to restrain the hydrogen bonds. The van der Waals force switch scheme with 1-1.2 nm cutoffs was used for the non-bonded interactions while the Particle Mesh Ewald (PME) summation with a radius of 1.2 nm was used for the electrostatics. The Verlet cutoff scheme was used for the neighbor’s list with a cutoff radius of 1.2 nm. Further details of the methods can be found in the SI.

### Laurdan Fluorescence

Laurdan dye was premixed with other lipids (molar ratio lipid:Laurdan: 200:1) and each AP in the initial organic solvent used to prepare the films. Two different lipid:AP molar ratios were used: 25:1 and 10:1. The details about sample preparation are found in SI. Fluorescence measurements were carried out by a Tecan Infinite 200 PRO microplate reader at 37 °C. The emission spectra of Laurdan were recorded between 390 and 600 nm, whereas the samples were excited at 355 nm. The generalized polarization (GP) at 355 nm excitation was calculated using fixed emission wavelengths of 440 nm and 490 nm according to Parasassi et al. (GP = (I_440_ − I_490_)/(I_440_ + I_490_)), where I_440_ and I_490_ are emission intensities at the characteristic wavelengths of the ordered phase (440 nm) and the disordered phase (490 nm). Laurdan GP provides information about the degree of hydration/lipid order in the polar head region near the glycerol backbone. Alterations in membrane water content and Laurdan dipole relaxation times cause changes in the excitation and emission spectrum of the probe. Lower GP values correspond to a more disordered lipid membrane (L_d_) while higher values to a more ordered membrane (L_o_ or L_β_).^52^

### Particle Size and Zeta Potential Measurements

The lipid vesicle’s size and zeta potential were measured at 37 °C with a Zetasizer Nano ZS analyzer (Malvern Instruments, Malvern, UK) equipped with a temperature-controlled cuvette holder. Two different lipid:AP molar ratios of 1:25 and 1:10 were used (additional info about lipid preparation found in SI). Approximately 1 mL of sample (200 μM final lipid concentration) was loaded into a square cuvette and ZEN1002 universal dip cell (Malvern) was placed in it. The dip cell stick is made of Teflon holding the Palladium electrodes. Dynamic light scattering (DLS) was used to determine the hydrodynamic diameter (Z-average, nm) of LUVs, calculated based on the intensity of scattered light. This measurement, a primary parameter provided by the instrument’s software, was derived using the Stokes-Einstein equation. The instrument has an attenuator available to adjust the laser power. Zetasizer determines the appropriate attenuator position during the measurement sequence. Zeta potential measurements, based on electrophoretic mobility, were conducted on the same sample to quantify the surface electric potential at the slipping plane using the Smoluchowski approximation, with a *f (κa)* value of 1.5. For accuracy, a minimum of three measurements were performed on each of three independently prepared samples.^53^

### Differential Scanning Calorimetry

Differential scanning calorimetry (DSC) experiments were performed on a Nano DSC microcalorimeter (TA instruments), using MLVs prepared as described above, in phosphate buffered saline (PBS), at different total lipid concentrations according to the lipid composition: 1.36 mM for DPPC; 3 mM for DPPC:SM (90:10), DPPC:Chol (95:5) and DPPC:SM:Chol (85:10:5). DPPC was used instead of POPC because the transition temperature of POPC (-4 °C) is not compatible with DSC measurements, PE and PS were not included as we decided to use simpler lipid model membranes. Drugs were co-solubilized with lipids in chloroform at a lipid:AP ratio of 10:1. A series of five consecutive DSC heating and cooling scans were performed using a scan rate of 1 °C/min and a temperature interval between 10 and 60 °C. Only the last scan, which was identical to the previous one, was analyzed. Data analysis was performed with the software Nano Analyze provided by TA instruments.

### Atomic Force Microscopy

For SUV preparation, the MLVs were sonicated over 3 cycles of 10 minutes, with an amplitude of 40 % and 3 s-pulses. SUVs were stored at 4 °C during a maximum 2-weeks. Supported lipid bilayers (SLB) were formed on freshly cleaved mica substrates by allowing the fusion of the SUV suspension (V = 100 µL, c = 1 mg/mL in buffer Tri-sHCl 50 mM, NaCl 500 mM, pH 8.0) at 60 °C for 30 min. Samples were then let for thermalization at ambient temperature for 15 min without dewetting and heavily rinsed with the appropriate buffer. When anionic lipids were used, the mica substrate was coated with 40 µL of CaCl2 10 mM during 10 min to favor the subsequent SUVs’ fusion onto the hydrophilic surface. Atomic force microscopy (AFM) imaging of the SLBs was then performed using the PeakForce Quantitative Nano-Mechanics (PF-QNM) mode on an Icon Fast-Scan setup (Bruker) in buffer conditions at room temperature (20 °C). Nitride-coated silicon cantilevers (ScanAsyst Fluid +, Bruker) with a nominal spring constant of ∼1 N/m were used and calibrated before each experiment using the thermal noise method. The images, analyzed and processed with the Nanoscope Analysis software (Bruker), were acquired with a scan rate of ∼1 Hz, a peakforce amplitude of 50 nm and a peakforce frequency of 1 kHz, where the applied force was kept as low as possible to minimize any damage (< 1 nN). Once the formation and stabilization of SLB has been confirmed by AFM imaging, samples were incubated with the AP of interest at a final concentration of 100 µM for 2 h. To probe the effect of APs, AFM images were finally recorded in different areas of the SLB. Along with the topographic images, AFM force curves were recorded in each pixel of the scanned area, and post-processed with Nanoscope Analysis using the Hertz model to estimate the local elasticity of the SLB, before and after AP treatment.

### Statistical Analysis

For ITC, particle size, zeta potential and Laurdan fluorescence measurements, data were analyzed using one-way ANOVA analysis. In case of ITC, the mean of each column was compared to the mean of all the other columns, and for the latter, the mean of each column (AP with lipids) was compared to the mean of control column (lipids alone). P-values smaller than 0.05 were considered significant (*), where values smaller than 0.01 were displayed with two stars (**) and the values smaller than 0.001 with three stars (***).

For AFM thickness and elasticity measurements, statistical significance was assessed using a two-sample t-test at a 0.05 level to compare conditions before and after incubation of APs with membranes.

## RESULTS

### CPZ and CLOZ have different membrane partitioning properties

To better understand the energetics of CPZ and CLOZ interaction with different lipid systems, and characterize it thermodynamically, isothermal titration calorimetry (ITC) has been used. The technique allows the determination of the partition constant (K_p_) between two media at equilibrium, in this case the AP partition between liposomes (LUVs) and aqueous phase according to the equation below:

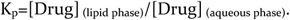

When the K_p_ values at pH 7.4 are compared with logP values (octanol/water partitioning), the same tendency is observed (Table 2). However, the use of liposomes, instead of octanol gives a more reliable value of the partitioning present in cells as the effect of phospholipid charges and other physico-chemical properties are intrinsically taken into account.^54^

Both APs possess quite high K_p_ with the three systems studied, as expected due to their high hydrophobicity. The value of K_p_ is higher for CPZ than CLOZ in all three systems, in agreement with the Marvin Sketch Simulations, predicting higher lipophilicity for CPZ as compared to CLOZ (Fig. S1). The obtained log*K*_p_ values for AP partitioning to POPC vesicles (system 2) are 4.12 ± 0.08 for CPZ and 3.49 ± 0.19 for CLOZ at pH 7.4, comparable to the ones obtained in the prediction of logP (5.41 and 3.23, respectively). Moreover, the *K*_p_ value characterizing the partition of CPZ partition to POPC obtained here (1.3 ± 0.2 x 10^4^) (Fig. 1a and Table S4) is in perfect agreement with the reported ITC value at the same pH (1.2 ± 0.4 x 10^4^).^24^ As for CLOZ, no ITC experiments have been found in the literature.

**Figure 1.**
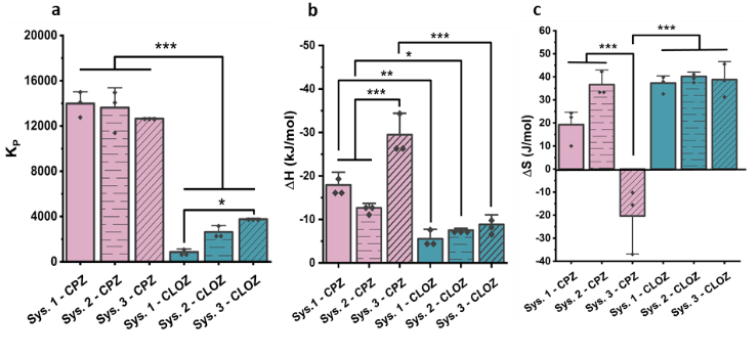
Thermodynamic parameters for the interaction of CPZ (pink) and CLOZ (cyan) with system 1, 2 and 3 at pH 7.4 obtained from ITC titration measurements. Panel a represents the partition coefficient, and panels b and c display enthalpy and entropy change, respectively. Three replicas of each condition are represented. One-way ANOVA test has been performed to compare each dataset between each other.

For both APs, adding anionic lipids (POPS) and curvature-promoting lipids (POPE) to POPC (system 3, Table 1) does not significantly impact the *K*_p_ (Fig. 1a and Table S4), also in agreement with reported data for CPZ.^19^ In the most complex system containing SM and Chol as well, no significant impact on *K*_p_ value has been observed for CPZ, as opposed to CLOZ, where *K*_p_ value decreases, which suggests that the lipid raft presence disfavors its partitioning. The enthalpy change (Δ*H*) is favorable in all systems and increased (in absolute values as it is negative) in presence of negative lipids (system 3) for CPZ (Fig. 1b and Table S4), which can be explained by the establishment of electrostatic interactions between the positively charged groups of CPZ and the carboxylic acid group and/or phosphate in DOPS, while the SM and Chol presence in system 3 likely reduces CPZ’s access to DOPS. The consistency of Δ*H* values for CLOZ across all three systems suggest that for this molecule, partitioning is more entropically driven. The enhanced electrostatic contribution for CPZ as compared to CLOZ, related with the higher increase in enthalpy variation, could be explained by the higher percentage of protonated CPZ molecules at pH 7.4 (Fig. S1). As for the entropy variation (Δ*S*), it is positive, thus favorable, for most systems, showing that the insertion into the bilayer of CPZ and CLOZ is entropy driven as well as enthalpy driven for most systems (Fig. 1c, Table S4). To recall, there are two major entropic contributions to the partition entropy that occur when APs partition from the water into the lipid membranes: an unfavorable contribution arising from the loss of degrees of freedom of the AP itself (called polarization entropy) and a favorable one arising from the release of water molecules that have been in contact with the AP once in solution (called solvation entropy). Additionally, a third contribution corresponding to the disordering of the lipids due to AP insertion should be considered. Total entropy variation is the sum of all these contributions.^55^ Indeed, the present study does not dissociate each of these entropy components, as we only obtain the overall entropy change. The only situation where an unfavorable entropy has been obtained is for CPZ interaction with system 3 (Fig. 1c). This suggests that the hydrophobic contribution to the partition of CPZ to the lipid bilayer is smaller for the charged membrane. This can be explained by the fact that enhanced electrostatic interactions occurring with charged membranes should result in CPZ location closer to the membrane surface .^56^ Indeed, electrostatic interactions often result in a negative entropy variation. An enthalpy/entropy compensation is common when the molecular association involves electrostatic interactions or other interactions that depend strongly on distance and/or orientation. Moreover, this CPZ sequestering by anionic lipids further decreases the degrees of freedom of this molecule once partitioned into the lipid system resulting in an increase in the unfavorable entropy contribution that is observed. CLOZ is interestingly not affected in the same way and its entropy contribution is the same or higher than CPZ’s entropic contribution, most probably due to its electrostatic interactions with the lipids being weaker (while both APs exist in an equilibrium between the non-protonated and protonated forms at the measured pH, the fraction of the charged form is higher for CPZ).

In addition to ITC, plasmon waveguide resonance (PWR) has been used to monitor AP affinity to membranes. PWR detects both mass and anisotropy changes in supported lipid bilayers upon addition of the APs (experimental details can be found in SI).^57^ Considering their very small mass, mass contributions to the spectra are negligible and spectral changes would result mostly from the reorganization of the lipid membrane in terms of anisotropy and mass density changes due to changes in lipid packing.^58^ Very small spectral changes have been observed by PWR upon incremental addition of both APs to the lipid systems, therefore it has not been possible to determine a binding affinity with this method. Nonetheless, since this is a surface binding method, the reversibility of the binding can be evaluated by following the spectral changes after washing the bilayer containing AP with buffer. While most CLOZ is removed upon washing, this is not the case for CPZ, suggesting stronger affinity and/or insertion depth of CPZ (data not shown).

ITC data provides evidence of AP partition to lipid membranes, with an important enthalpy contribution, which in turn indicates that, in addition to entropic contribution, these interactions are also driven by electrostatic forces. To quantify the changes in electrostatic drug-lipid interactions, the zeta potential values of liposomes prior and after AP interactions have been evaluated, where data interpretation considered that the AP molecules are charged.^59^ Quasi neutral zeta potential value (∼ 0 mV) has been measured for system 2, whereas negative values (-13 and -16 mV, respectively) have been found for system 1 and 3, in line with the presence of the negatively charged lipid DOPS (Fig. S3). Overall, the addition of both APs results in an increase in zeta potential values, making them less negative, and this effect increases with AP concentrations as a result of the charge neutralization brought by the protonated AP molecules. These results underscore the importance of electrostatic interactions in the mechanism of AP membrane affinity. Both approaches, ITC and zeta potential measurements provide complementary insights into the AP binding and bilayer charge alteration, which further elucidate the nature of the drug-lipid interactions. The obtained results are consistent with data from the scientific literature on CPZ.^60^

### Both APs locate at the phosphate/glycerol level

To further characterize the distribution and location of the two AP molecules, all-atom molecular dynamics (AAMD) simulations have been used (Fig. 2) with non-charged APs. Consistent with the ITC measurements, both APs have been shown to interact with the membrane for the three lipid systems investigated. Simulations additionally show that APs insert near the lipid headgroup region at the phosphate or glycerol level. Some differences are observed between the two APs, the lipid systems and the lipid:AP ratios investigated as summarized below.

**Figure 2.**
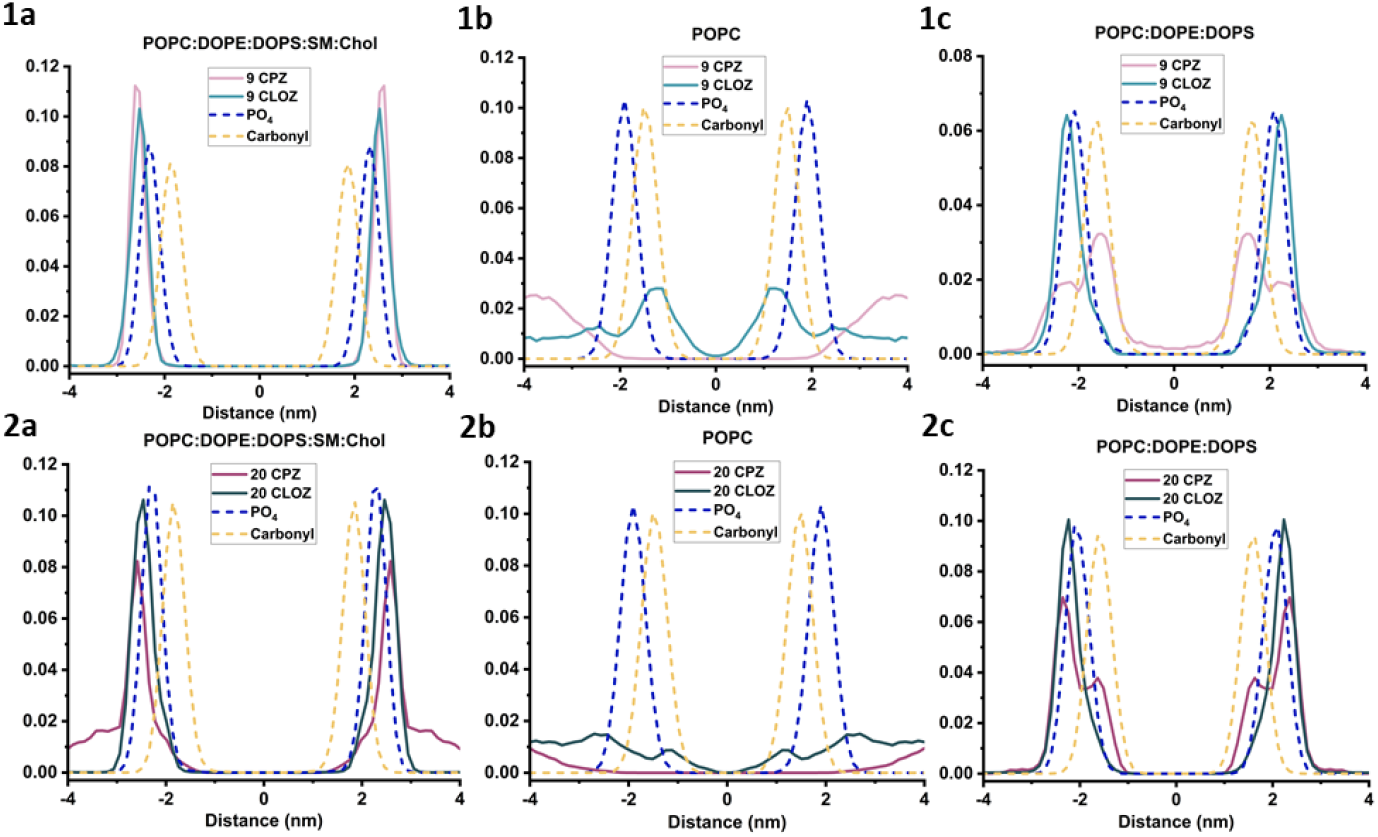
Normalized relative density of the AP chlorine atoms and membrane phosphate and carbonyl groups in: system 1 (1a and 2a), system 2 (1b and 2b) and system 3 (1c and 2c) determined by AA-MD simulations. The panels represent the density in the presence of 9 AP molecules (1) and 20 AP molecules (2) (either CPZ - pink or CLOZ - cyan).

In the three systems, it has been found that both CPZ and CLOZ tend to form clusters in water, with CPZ forming larger aggregates than CLOZ. This likely results from the higher hydrophobicity of CPZ (logP(CPZ) = 5.41, logP(CLOZ) = 3.23) which hence tries to reduce its surface area in contact with water. Such clustering often leads to reduced interactions with the membrane.

In system 1, reflecting SVs, single density peaks are observed for both CPZ and CLOZ (Fig. 2, 1a and 2a) that are localized just above the phosphate groups. Both molecules interact with the membrane, although these interactions are mostly at the shallow water-membrane interface and not deeper than the phosphate groups. CPZ forms larger aggregates than CLOZ, but despite that still penetrates and inserts in the membrane near the phosphate group region. In the more simplistic system 2, and unlike in system 1, CPZ does not penetrate into the membrane while CLOZ penetrates deeper than in system 1, around the glycerol group level (Fig. 2, 1b and 2b). Previous modelling studies on the same system (POPC) have revealed the non-protonated form of CLOZ to locate around the phosphate group when at low molar ratios.^61^ CPZ forms large aggregates which do not embed in the membrane at any point and instead remains solvated in water. Such observations suggest that the presence of negative lipids in system 1 facilitates the interaction between CPZ and the lipid bilayer, which is not a prerequisite in the case of CLOZ. Since system 1 also contains SM and Chol, their contribution to AP interaction has been also investigated in system 3. CPZ, despite forming larger aggregates, penetrates deeper into the system 3 membrane than CLOZ. CPZ also presents a double peak at 1.5 nm and 2.5 nm distance away from the bilayer center. The inner peak (1.5 nm) correlates with the glycerol groups, thus the CPZ molecules are embedded deep in the membrane while the outer peak (2.5 nm) lies just above the phosphate level. For CLOZ, only one density peak is observed above the functional groups. Despite CLOZ forming smaller aggregates than CPZ, it penetrates system 3 less deep than CPZ. Such behavior is consistent with the higher lipophilicity of CPZ.^62^ The ability of CLOZ to embed deeper in system 2 unlike in systems 1 or 3 is probably due to the absence of DOPE/DOPS headgroups since the packing and area per lipid are similar in system 2 and 3. In the system 3, a different insertion distribution has been observed at different CPZ ratios: when lower proportion of CPZ (9) are used, most ligands are inserted deeper in the membrane (Fig. 2, 1c), while the opposite is seen for the system with 20 CPZ, due to aggregation (Fig. 2, 2c).

Overall, both APs locate mostly near the phosphate/glycerol level. The preference for the lipid headgroup of the aromatic moieties is largely known to arise from dipole interactions between the aromatic groups and lipid headgroups.^63^

Electrostatic interactions impact the affinity of both APs for the lipid membrane, especially for CPZ, as seen in ITC. Therefore, in presence of anionic lipids, the APs are sequestered at the phosphate/glycerol and CPZ molecules are found deeper in the membrane (Fig. 2, 1c and 2c). In the complex lipid systems, preferential contacts of the APs are observed with Chol and SM. This is in good agreement with the fact that APs have been shown to preferentially partition into the so-called raft domains in membranes.^64^

### The impact of APs on membrane’s physical and mechanical properties

After revealing that both APs partition in different lipid membrane systems in absence or presence of anionic lipids as well as SM and Chol, their impact on membrane’s physico-chemical and mechanical properties has been investigated, namely membrane order, area per lipid, phase transition, thickness, elasticity, integrity (pore formation), liposome size and charge using a multi-approach strategy.

### Both APs increase membrane order parameter of disordered membranes

The impact of APs on the membrane order parameter along the phospholipids’ fatty acid chains has been investigated by AA-MD simulations. In addition, to access the lipid headgroup region, fluorescence spectroscopy has been used by incorporating Laurdan, a fluorescent reporter dye sensitive to dipolar relaxation processes and the polarity of the microenvironment in the polar headgroup region near the glycerol.^52^ Therefore, these complementary approaches allow assessing the role of APs in both headgroup and fatty acid chain regions of the lipid membrane.

System 1 represents a very compact membrane as seen by its area per lipid being the lowest of the three systems (Table S1). The extended lipid chains increase thickness and decrease the area per lipid (Fig. S4, Table S1). For such a membrane the order parameters (Scd) are high as seen in Fig. 3. A general increase in order parameter is observed for all lipid types, POPC, DOPS and DOPE in presence of both APs at low and high concentrations (Fig. 3). Correlated with the order parameter, the area per lipid has also been analyzed by AA-MD simulations, where both APs lead to reduction of the area per lipid, especially at lipid:AP ratio 100:9 (Table S1), and an increase in membrane thickness (Fig. S4). This is in accordance with the higher order parameter induced by APs, in general inducing a tighter packing and reduced area per lipid. The Scd value for POPC in the presence of 9 CLOZ is the highest, followed by 9 CPZ which is higher than the pure membrane system. This indicates that CLOZ is able to embed better in the membrane than CPZ and increase the lipid chain stiffness as noted by the higher order parameter (Fig. 3).

**Figure 3.**
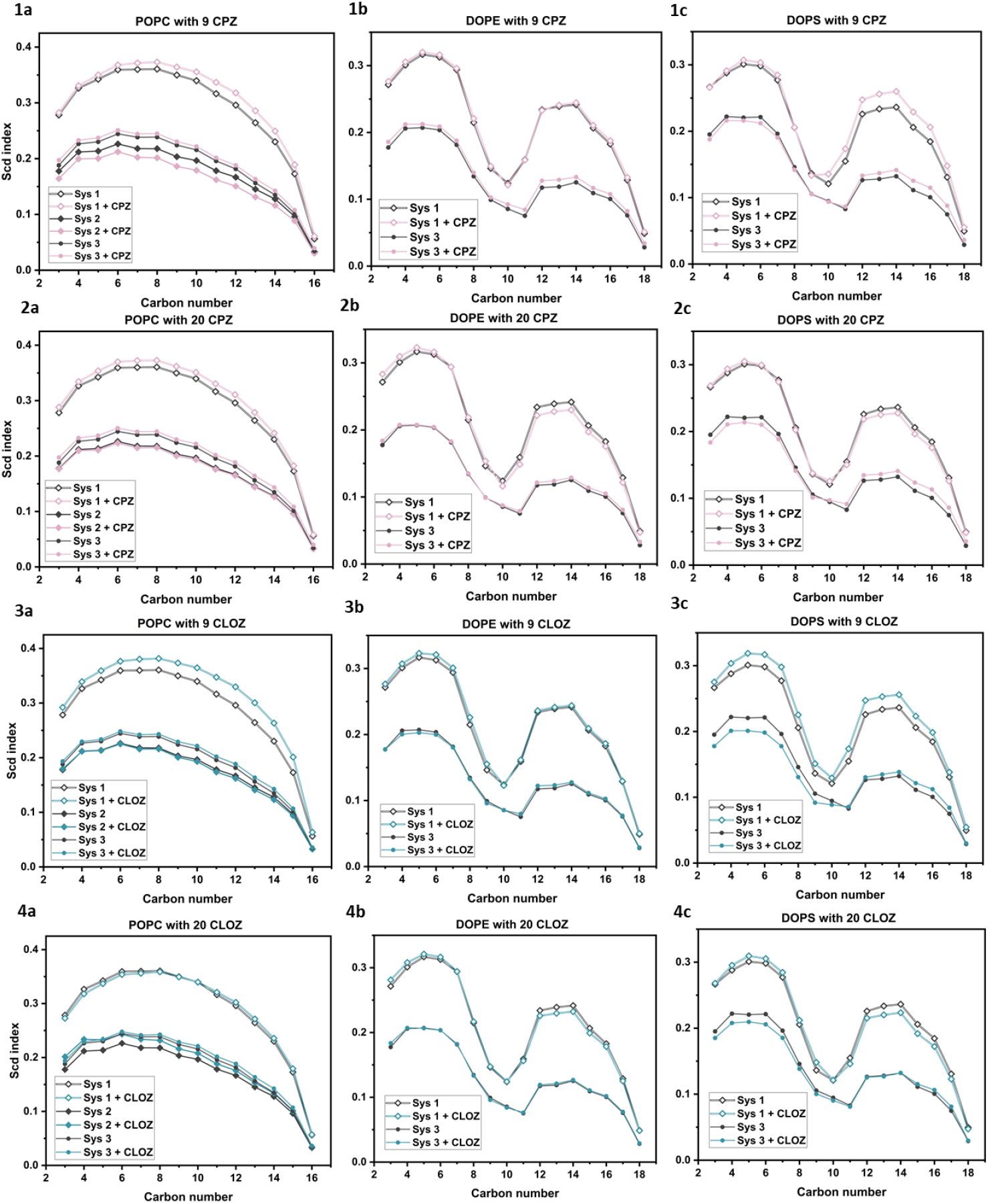
Effect of CPZ (pink) and CLOZ (cyan) on lipid chain Scd (sn1) in systems 1-3 obtained by AA-MD simulations. The effect of CPZ with 9 (1) and 20 (2) molecules is shown on different phospholipids: POPC (1a, 2a), DOPE (1b, 2b) and DOPS (1c, 2c). The effect of CLOZ with 9 (3) and 20 (4) molecules on different phospholipids: POPC (3a, 4a), DOPE (3b, 4b) and DOPS (3c, 4c) is represented.

In system 2, membrane thickness is lower in presence of CPZ, whereas it is about the same or higher for CLOZ when compared to POPC alone (Fig. S4). This indicates that, especially at lipid:AP ratio 100:9, the CPZ molecule presence in the membrane displaces the lipids that move further apart, increasing the area per lipid and reducing the thickness (Table S1 and Fig. S4). The effect observed shows the opposite trend than the system 1, which may be due to system 2 being less compact and with a reduced thickness, which facilitates the lipid displacement observed for CPZ.

In the case of system 3, a slight change is observed in the DOPS chains while POPC and DOPE chains show almost no change with any of the two APs (Fig. 3). This might be due to the larger area per lipid in system 3 compared to system 1 (Table S1). System 1 is tightly packed due to the presence of Chol and thus has higher Scd and thickness. System 2, however, is sparsely packed and represents a homogenous mixture of a pure lipid. System 3 introduces more lipid species that can form favorable interactions with the APs in the absence of SM and Chol. As such it provides a more favorable environment to accommodate CPZ as is observed by the density profiles in Fig. 2, 1c and 2c, and the corresponding reduction in membrane thickness (Fig. S4) and higher area per lipid (Table S1). In CLOZ presence, reduction in membrane thickness (Fig. S4) and area per lipid (Table S1) have been observed while the relative density profile has remained similar to the one observed in system 1 (Fig. 2, 1a, 2a, 1c, 2c).

Overall, AP impact on order parameter is highly dependent on the lipid:AP ratios and the results with 20 AP molecules are quite different due to the aggregation tendency of APs hampering their membrane insertion. This observation is in good line with reported aggregation of CPZ.^60^ The CPZ molecules tend to clump and form large aggregates while CLOZ forms smaller aggregates. At high lipid:AP ratio, more APs are part of an aggregate and only a small fraction of that aggregate interacts with the membrane, hence the lowered effect of the APs on the order parameter, membrane thickness and area per lipid.

To summarize the main findings of the effect of APs on fatty acid chain ordering across the studied lipid mixtures: a general increase in the molecular order parameter is found, suggesting a decrease in membrane fluidity upon ligand incorporation. Prominent changes for POPC order parameter in the three systems are observed while minor changes are seen for DOPE and DOPS. However, this effect is minimized in the simplified system 2 and 3, which could be due to the absence of Chol and SM. Indeed, the strongest impact in ordering has been seen in POPC and DOPS in system 1. This trend of order parameter increase could also change depending on the lipid:AP ratio. Higher AP amounts can have a lower effect on the order parameter for both APs due to their propensity to form large aggregates (more often observed with CPZ). The distinct interactions between CPZ and CLOZ with the lipid bilayer probably originate from the unique localization and orientation of CLOZ close to glycerol backbone compared to CPZ found rather at the phosphate group. This differential interaction arises from the variations in the molecular structure and polarity between CPZ and CLOZ.

Laurdan fluorescence has been measured for the 3 lipid systems in the absence and presence of both APs at lipid:AP ratios 25:1 and 10:1. Fluorescence emission spectra of Laurdan-labelled lipid mixtures with and without APs are presented in Figure 4. The emission spectrum in POPC alone (system 2) shows two peaks, centered at 435 nm (blue-shifted, ordered environment) and 490 nm (redshifted, disordered environment) (Fig. 4, 2b). The spectra of AP-containing lipid mixtures also exhibited these two peaks with the intensity of the first peak increasing as a function of AP concentration and type of the ligand used. The two detected peaks indicate the coexistence of two distinct polarity microenvironments sensed by the Laurdan probe. This can be attributed to the presence of palmitoyl (saturated) and oleic (unsaturated) fatty acid chains in the POPC molecules, suggesting distinct polarity of Laurdan within the lipid bilayer. It is noteworthy to mention that even DOPC or DLPC membranes in liquid-disordered phase exhibit two peaks Laurdan emission spectra at physiological temperature.^52,65^ It is suggested that Laurdan resides in membrane sites (cavities) with varying water content, affecting its GP value due to differences in dynamically restricted water molecules.^66,67^ As the ligands influence the blue-shifted peak, it suggests that both APs primarily interact with the ordered environment and reduce the polarity sensed by Laurdan, hence decrease the access of water to Laurdan. The opposite trend has been observed with systems 1 and 3, where AP presence increased the polarity sensed by Laurdan, demonstrated interaction with the disordered environment by variation in the intensity of red-shifted peak (Fig. 4, 2a and 2c). The calculated Laurdan GP of the three studied lipid bilayers follows this membrane lipid order rank: system 2 < system 3 < system 1 (Fig. 4, 1a-1c). Both APs dehydrate the membranes, which mostly goes in the direction of the increase the membrane lipid order at the glycerol level, generally observed (Fig. 3). CLOZ demonstrates higher potential to change the lipid order than CPZ. CLOZ induces changes in lipid order in the following rank: system 1 < system 3 < system 2. System 1, containing SM and Chol, represents the membrane model with the highest lipid order among the studied systems. The ordering effect of CLOZ has been constrained by the already highly ordered structure of this system.

**Figure 4.**
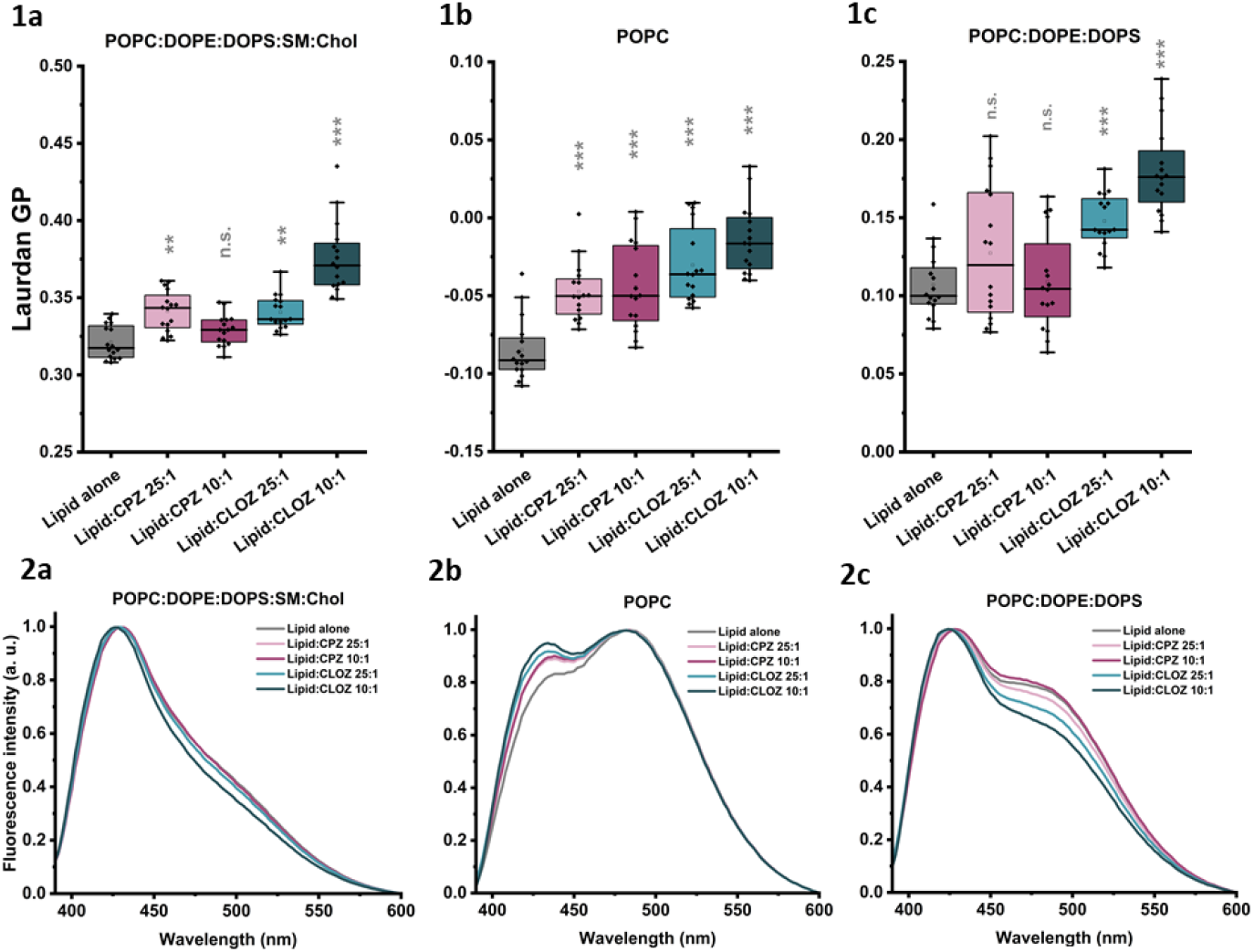
Effect of APs on the lipid order of LUVs as measured by Laurdan fluorescence spectroscopy. Laurdan GP values (1) and fluorescence intensity spectra (2) of control LUVs without AP (grey) and AP-containing ones in lipid:AP ratios 25:1 and 10:1, where CPZ is represented in pink and CLOZ in cyan. Three different lipid systems are represented: system 1 (a), system 2 (b) system 3 (c). The measures have been performed at 37 °C. One-way ANOVA test has been performed to compare each set of data with the control group (lipid alone, test results present above the data populations).

Thus overall, the Laurdan data points to an increase in GP which correlates with an increase lipid ordering in the headgroup region that is more pronounced for CLOZ than CPZ. In view of the fact AA-MD simulations show that the molecules are located around the headgroup region, their accumulation at this level decreases the relaxation rates of the water dipoles as well as Laurdan excited state dipole moment which could result in an increase of ordering.

One issue worth considering relates to the fact that the AP effect on membranes is highly dependent on the lipid composition. Indeed, the AP-induced lipid ordering observed in this study contrasts with the disordering effect detected by fluorescent probes in studies involving dopamine antagonists in natural lipid extracts.^68^ The lipid composition of the membrane is therefore a crucial *factor* influencing AP effects on lipid order. In line with such observations, then we ask the question whether the changes in membrane order parameter induced by APs depend on the inherent membrane ordering state, so we investigated AP impact on the phase transition of saturated and thus ordered phospholipids.

### APs stabilize the fluid phase of ordered lipid membranes

The phase transition temperature of lipids is a property that is quite sensitive to the presence of foreign molecules and can provide indirect information about their mode of interaction (insertion level, homogeneity in distribution, etc.) and effect on membrane properties (ordering, membrane curvature). Two approaches have been used to evaluate AP impact on lipid phase transition: differential scanning calorimetry (DSC) and solid state nuclear magnetic resonance (ssNMR).

The effect of both APs on the phase transitions of lipid bilayers composed of DPPC, DPPC:SM (90:10), DPPC:Chol (95:5) and DPPC:SM:Chol (85:10:5) has been assessed using DSC. DPPC has been chosen as its fatty acid chain length (C16) is comparable to that of POPC (C16, C18) and its melting temperature (T_m_) is 41 °C, so well above 0 °C (required for DSC studies).^69^ When added at a lipid:AP molar ratio of 10:1, both APs lead to a shift of the main transition peak towards lower temperatures, for all the considered lipid systems, suggesting that they facilitate the transition to the fluid phase (Fig. 5). Also, the presence of the APs leads to a broadening and flattening of the transition peak, while not changing significantly the main transition enthalpy. Interestingly, the largest decrease in transition enthalpy occurs for CPZ with DPPC/SM/Chol. Overall, the effects observed with CPZ are always stronger than with CLOZ, suggesting its better insertion in the bilayer, which goes in line with ITC partitioning. It should be noted that the effects observed with CLOZ are consistent with those previously published in the literature.^20,70^ The overall decrease in the gel to fluid phase transition temperature shows that these APs interact preferentially with the fluid phase. This in turn indicates that they will exert a disordering effect in the membrane.^71^ In line with the results, a recent study on a model membrane composed of DPPC, SM and Chol, using DSC at three pH values and as a function of drug concentration of seven APs including CLOZ, also showed that APs induce more fluidity and disorder in such lipid systems.^70^ Palmeira et al. found a direct correlation between the membrane disordering effect of APs and their *K*_p_. The higher *K*_p_ lead to higher local concentrations in the membrane and thus a higher effect in the main transition.^68^ While we have not investigated a sufficient number of AP molecules to draw a definitive conclusion, it is noteworthy that the more significant decrease in lipid phase transition temperature (Table 3) is observed for CPZ, which has a higher K_p_ than CLOZ, supporting this correlation.

**Table 3.**
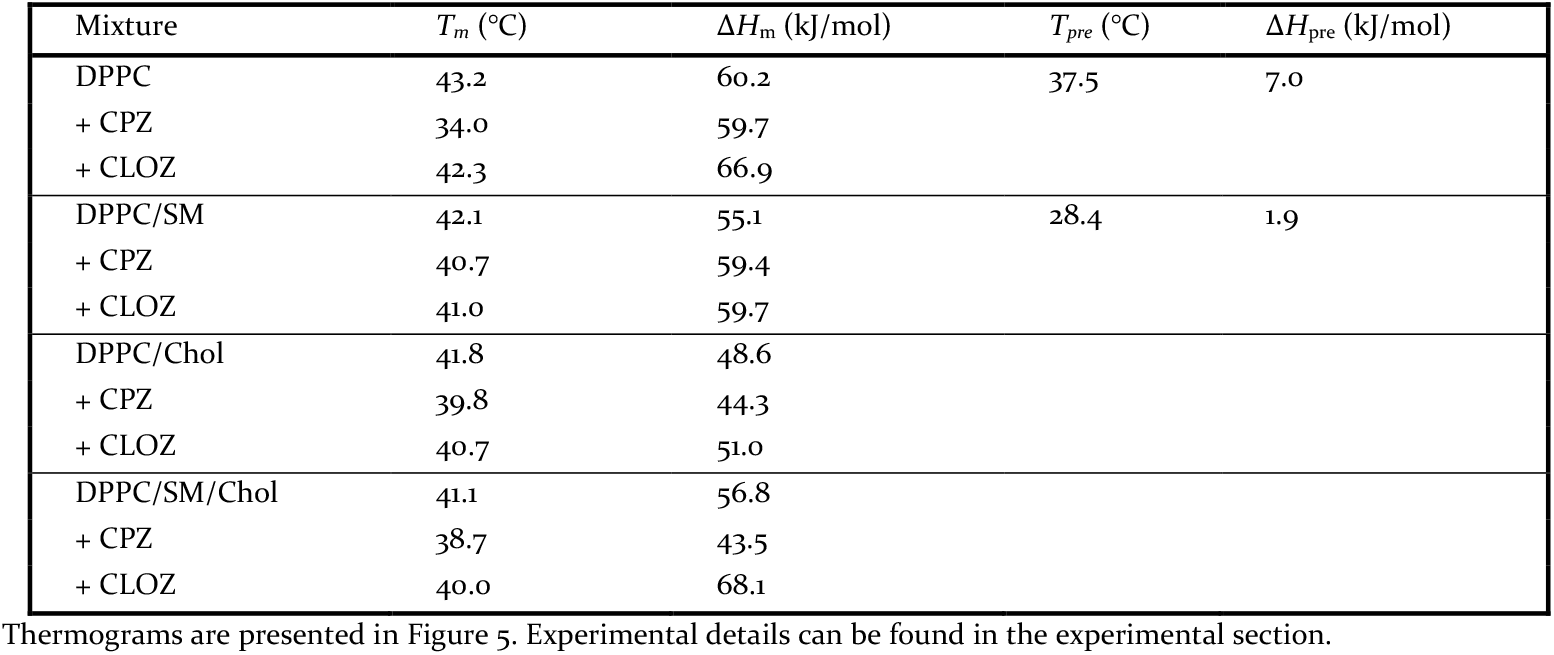
Thermodynamic parameters of the lipid phase transitions obtained by DSC for the lipid systems (DPPC; DPPC/SM 90:10; DPPC:Chol 95:5 and DPPC:SM:Chol 85:10:5) in the absence and presence of lipid:AP ratio 10:1. Each mixture was measured one time. Thermograms are presented in Figure 5. Experimental details can be found in the experimental section.

**Figure 5.**
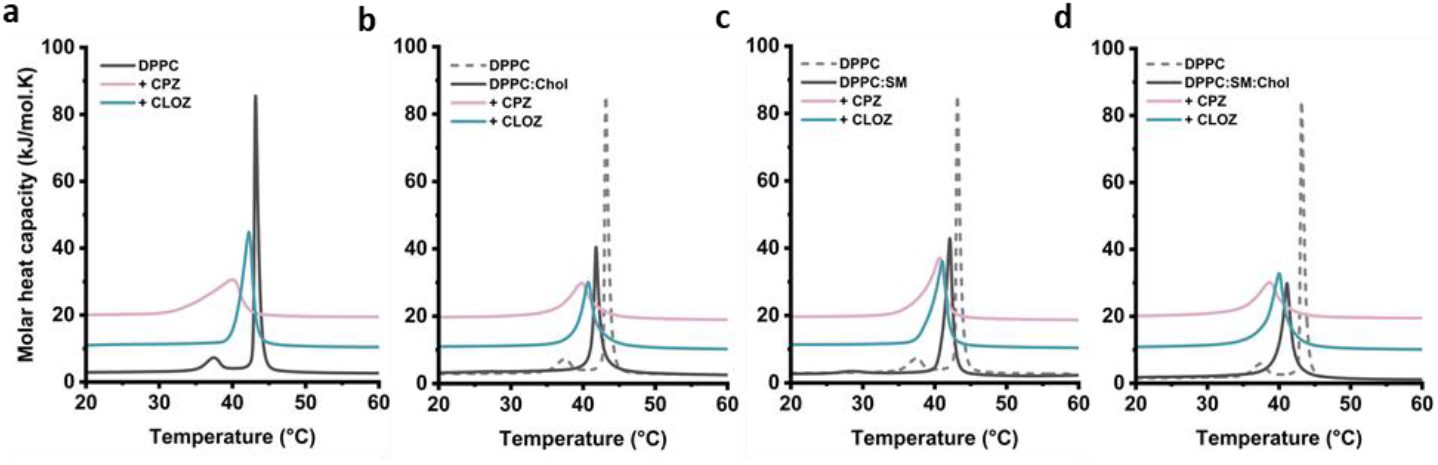
Representative DSC heating thermograms. DPPC (a), DPPC:Chol (50:5, b), DPPC:SM (90:10, c) and DPPC:SM:Chol (85:10:5, d) systems are measured alone (grey) or in the presence of CPZ (pink) or CLOZ (cyan) at an lipid:AP molar ratio of 10:1.

In the case of fluid membranes investigated by AA-MD simulations, an increase in the order parameter has been induced by APs. DSC data shows a preferential interaction of the APs with the fluid phase, thus suggesting a membrane disordering effect. It seems that the effect of AP on membrane ordering is highly dependent on the lipid state *per se*, with an increase in membrane ordering when starting in the fluid state and a decrease in membrane ordering when initially in the gel state. To test this hypothesis ssNMR has been performed.

To make the parallel with the DSC experiments, M1 profiles by ssNMR have been acquired using the same lipid – DPPC for one of the APs. CLOZ impact has been the opposite depending on the state of the bilayer with a disordering effect observed below the phase transition (in the gel phase, CLOZ curve is below DPPC) and an ordering one above the phase transition (in the fluid phase, CLOZ curve is above DPPC) (Fig. S5). This dual effect can explain the differences observed with AA-MD simulations: while by DSC, the addition of APs to a more ordered DPPC provokes a disordering effect; with AA-MD simulations, AP addition has the opposite ordering effect on the membranes that are initially more disordered as POPC is used with lower T_m_, suggesting that APs might exert a Chol-like effect on membranes.

### APs impact membrane thickness and elasticity without affecting membrane morphology

Membrane ordering being correlated to membrane thickness, the AP impact on membrane thickness has been investigated by AA-MD simulations and AFM. In addition, AP impact on the nanoscale morphological (formation of holes or other defects) and mechanical properties of supported lipid bilayers (SLB) such as elasticity has been assessed by AFM. Both pure POPC membranes and more complex ones containing SM and Chol do not reveal any phase separation, and are deposited as homogeneous and flat bilayers (Fig. 6a). Defects could be found over the scanned areas, and served as a “visualization control” to estimate the bilayer thickness (Fig. 6b). Incubation with either CPZ or CLOZ at a concentration of 100 µM for 2 h has not resulted in any apparent alteration (aggregates, holes, etc.) of the lipid bilayer (Fig. 6a), consistent with their mechanism of action, opposed to *e*.*g*. antimicrobial peptides that would disrupt and destroy the SLB over time.

**Figure 6.**
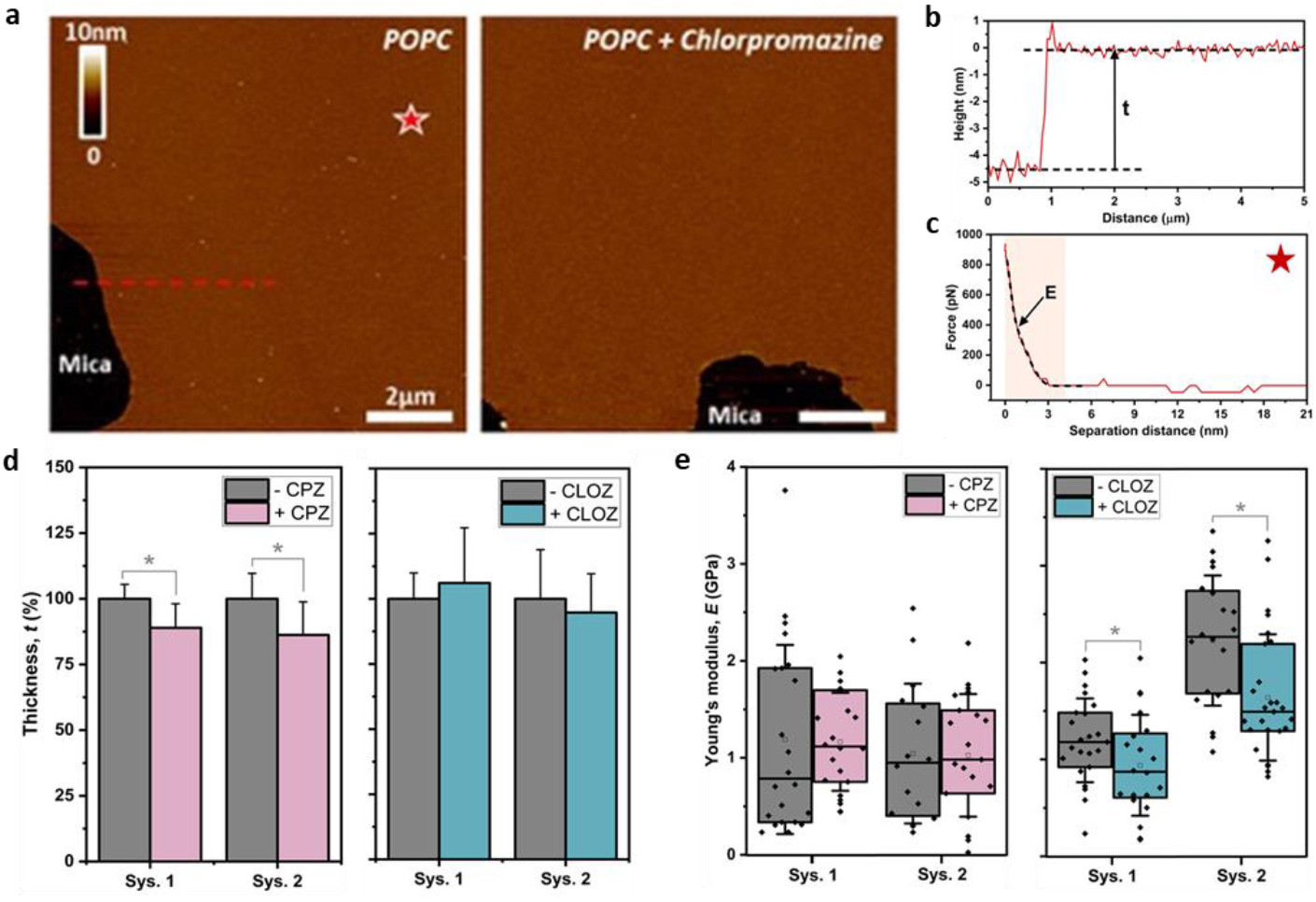
Nanoscale investigation of AP impact on lipid membranes. (a) Representative topographic AFM images of a POPC bilayer before (left) and after (right) injection of 100 µM of CPZ, on two distinct areas. (b) Height profile along the dashed line in (a) allowing the estimation of the bilayer thickness (t). (c) Representative force curve recorded on the POPC bilayer (red star in a) allowing the estimation of the apparent young’s modulus (E) of the bilayer. (d) Influence of CPZ (pink) and CLOZ (cyan) on the bilayer thickness t (here normalized and expressed as %) depending on the lipid composition (system 1 and 2), as determined from the AFM images and subsequent height profiles. (e) Influence of CPZ and CLOZ (pink and cyan, respectively) on the apparent young’s modulus of the bilayer. The error bars stand from the standard deviation are estimated from at least 3 independent replicates.

The potential impact of APs on membrane integrity and potential pore formation has also been investigated by a dye leakage assay that consists in encapsulating a dye (sulforhodamine) in liposomes at concentrations that result in auto quenching. The possible destabilization of the membrane integrity by APs results in dye leakage and a consequent increase in fluorescence intensity. Overall, the two APs do not perturb the membrane integrity following 1 h incubation, except for system 3 where CPZ but not CLOZ shows a concentration-dependent impact (about 10 % leakage maximum reached at lipid:AP ratio of 10:1) on membrane integrity (Fig. S6). The increase in leakage promoted by CPZ in system 3 might be the result of phase separation being induced by this AP in such system, although this needs further investigation. Dose-dependent perturbation of membrane integrity by CPZ has been previously reported in the literature with up to 100 % leakage in liposomes (POPC; POPC:SM:Chol 3:1:1). Their study, however, displays much higher lipid/AP ratios with similar lipid concentration, whereas our study has focused on the maximal lipid:AP ratio of 5:1.^72,73^

This observation is in line with the AFM results described below where CPZ tends to induce membrane thinning in contrast to CLOZ. The difference regarding both techniques (leakage assay and AFM) may result from the different experimental conditions (planar membranes versus liposomes, lipid:AP ratios and incubation time).

The use of AFM spectroscopy for measuring the membrane thickness (Fig. 6b) has not displayed morphological damages described above. CPZ tends to induce a 10-15 % thinning of the SLB, whatever the composition (from 5.2 to 4.6 nm for system 1 and from 4.8 to 4.1 nm in case of system 2) (Fig. 6d). While AFM reveals a decrease in membrane thickness by CPZ in both systems 1 and 2, AA-MD simulations reveal a general increase in thickness for system 1 (Fig. S4) after the addition of CPZ, which is in accordance with reduced area per lipid and more dense packing of the bilayer observed (Table S1). This correlates with the general increase in Scd (Fig. 3). In the case of CLOZ, a similar thickness increase has been observed by AA-MD simulations, while no changes have been observed by AFM. For system 2, like in AFM, membrane thinning has been observed by AA-MD simulations for CPZ, especially 9 CPZ, since in presence of 20 CPZ aggregate formation reduces the insertion of the molecules within the bilayer. The corresponding reduction in POPC Scd for system 2 with 9 CPZ has been observed (Fig 3, 1a). In case of CLOZ, no changes in membrane thickness have been observed by AFM, while in AA-MD simulations thickness increase has been observed at high ratio (20) that corresponds to the increase in POPC Scd in Fig. 3-2a. In system 3, there is a general thickness decrease observed for both APs by AA-MD simulations but interestingly, only a reduction in the DOPS Scd and not for POPC, DOPE has been observed (Fig. 3). The thickness differences observed between AFM and AAMD simulations can be explained by the shorter timescale studied in AA-MD simulations and initial drug aggregation in water, as well as by the use of SLB in AFM studies.

AFM force spectroscopy has also been used to probe the elasticity of the bilayer: Young’s modulus has been estimated from the Hertz model. Surprisingly, the thinning effect observed for CPZ is not associated with any mechanical alteration of the SLB, as illustrated by similar Young’s modulus of the SLB before and after CPZ action (Fig. 6e). This might suggest a subtle change in the lateral organization of the lipid membrane when CPZ interacts with the bilayer without significant variation in the whole membrane elasticity. Interestingly, CLOZ shows an “invert” behavior: while it does not impact the bilayer thickness, it induces a decrease in its Young’s modulus, suggesting a higher elasticity, and potentially fluidity, of the membrane, whatever its composition.

### Both APs decrease liposome size

The DLS method has been used to monitor the effect of APs on the size distribution of LUVs in lipid systems 1, 2 and 3 (Fig. 7). For all three studied systems without APs, narrow monomodal size distributions centered between 131 and 139 nm are obtained (Fig. 7 and Table S5). The vesicle size in system 3 decreases compared to system 2, which can be attributed to the presence of PE and PS. In contrast, the further addition of SM and Chol in system 1 increases vesicle size, although it does not exceed that of POPC. Both APs, within the studied concentration range, reduce LUV sizes by up to 5 % across all three systems. This reduction in size may result from the overall membrane ordering effect of the APs in the phospholipid chain region, leading to a decrease in the area per lipid.

**Figure 7.**
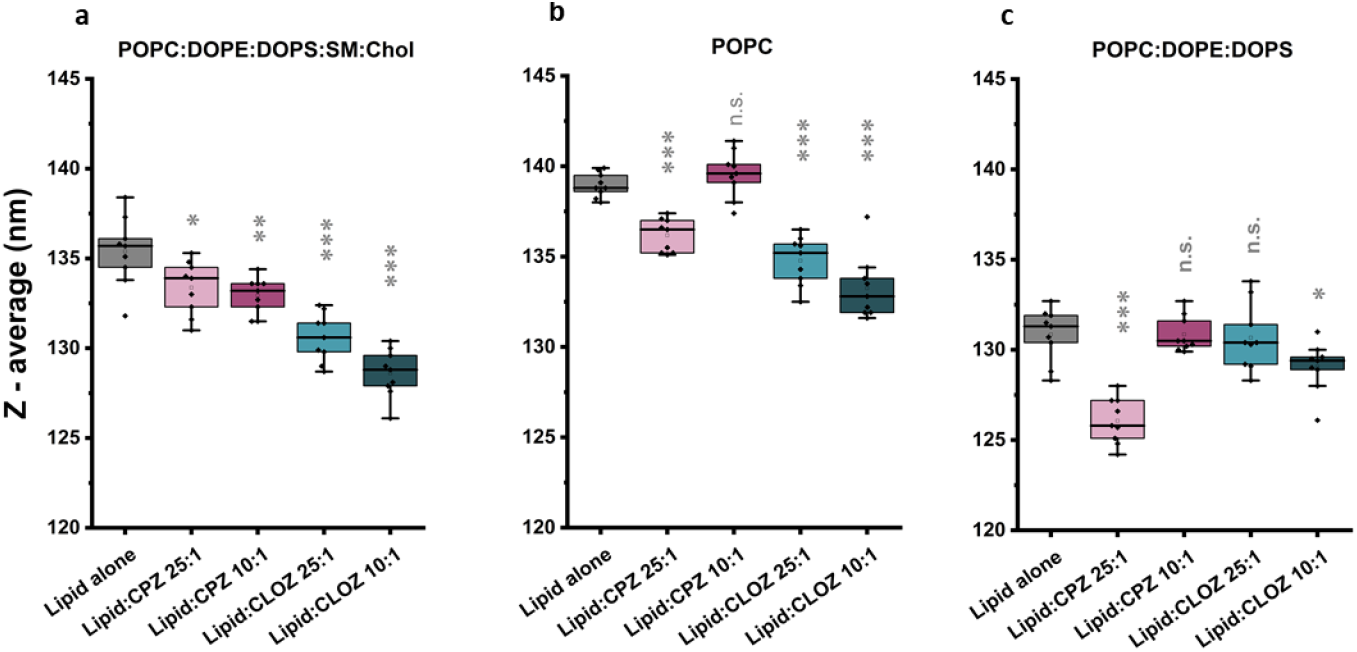
Z-average size of control LUVs without AP (grey) and AP-containing ones (CPZ – pink, CLOZ - cyan) in lipid:AP ratios 25:1 and 10:1. Three different lipid compositions have been used: system 1 (a), system 2 (b) and system 3 (c). One way ANOVA test has been used for comparing each dataset with the control group (lipid alone, the results are shown above each box).

In brief, two distinct modes of interaction between the APs and lipid membranes are observed. CLOZ consistently reduces vesicle size, with the smallest vesicle size observed at the highest CLOZ concentration. The size reduction following CLOZ addition shows a similar trend in systems 1 and 2, with the most significant reduction (5 %) observed in system 1. Both CPZ ratios reduce vesicle size in system 1.

In contrast, in systems 2 and 3, CPZ initially decreases vesicle size at a lower lipid:AP ratio (25:1). However, at higher ratios, CPZ leads to an increase in vesicle size, with values generally returning to those observed in the control condition. Notably, CPZ addition results in the greatest reduction in vesicle size within system 3.

## CONCLUSIONS

APs are known to exert their clinical activity primarily by modulating neurotransmitter systems, particularly dopamine pathways, by an antagonistic competition with the endogenous ligands at different subtypes of brain G-protein coupled receptors (GPCRs). However, their interaction with membranes affects their apparent receptor affinity and the biophysical changes observed may affect the signaling cascades.^13,14^ Therefore, this study focused on the analysis of AP-membrane interaction.

Regarding the characterization of CLOZ and CPZ membrane interactions, seminal work by Jutila et al. on MLVs of mixtures of DPPC and brain PS, using DSC and Langmuir balance, showed that CLOZ could abolish phase separation in the mixture and induce a greater increase in the average area/molecule as compared to CPZ.^20^ Consistent with Jutila’s results that show a strong partitioning into brain PS for both APs, our results show that CPZ has a higher partition coefficient (*K*_p_) than CLOZ in all lipid systems, regardless of the differences in lipid composition. The interactions, as expected, have been stronger for CPZ as compared to CLOZ (where the interaction is more entropically driven), a result that we explain by the fact that CPZ is mostly protonated at physiological pH while CLOZ is present in both protonated and non-protonated forms.

Based on AA-MD simulations (non-protonated form), in system 1 both CPZ and CLOZ are predicted to be located just above the phosphate groups of the membrane while some CPZ molecules have been observed at a deeper level, especially at low lipid:AP ratios (Fig. S7). This goes along with reported literature on CPZ, showing that it is positioned with the aromatic rings just below the glycerol level with one of the rings establishing van der Waals interactions with the methylene groups of the first carbons in the fatty acid chain.^27^ Although CPZ tends to form larger aggregates than CLOZ, it partitions better to the lipid model membranes. However, CLOZ can be found both at the phosphate groups and at the glycerol level.

Regarding the impact of APs on membrane order parameter, the results are quite surprising as they are highly dependent on the inherent membrane ordering nature. For the disordered lipid membrane systems investigated (system 1, 2 and 3) both the addition of CPZ and CLOZ resulted in an increase in the order parameter, with the magnitude of the effect being much greater for CLOZ, particularly in system 1 which is the less disordered suggesting the possibility of more ordered synaptic vesicle membranes after CLOZ treatment. Conversely, the AP impact on highly ordered lipids (like DPPC) was found to be entirely different, as both APs have been shown to preferentially interact with the fluid phase. The decrease in the enthalpy of the lipid transition by the APs is larger for CPZ than CLOZ, which correlates with the observation that some CPZ molecules are often found deeper in the membrane than CLOZ. This dual behavior of APs depending on the membrane gel/fluid state recalls that observed by Chol presence in lipid membranes and has been observed for other APs, suggesting a Chol-like behavior of APs within lipid membranes.^12^

Moreover, preferential interactions of both APs with Chol have been observed, probably as a result of favorable π-interactions between both molecules. AFM evidenced that while the addition of CPZ induced membrane thinning of the bilayer in systems 1 and 2, surprisingly this effect did not result in variation of the mechanical properties of the membranes. In contrast, although the addition of CLOZ did not affect the bilayer thickness, it did decrease the Young’s modulus, indicating a higher elasticity and possibly fluidity of the membranes despite a greater order parameter.

Vesicle size reduction has been observed with both APs, with the greatest reduction (5 %) being observed after CLOZ addition in system 2, consistent with greater dehydration caused by CLOZ compared to CPZ, in line with the location of CLOZ at the glycerol backbone or glycerol/phosphate level. Overall, the addition of both APs resulted in an increase in zeta potential values, making them less negative. The results from system 1 suggest a modulatory role of Chol in the localization of CLOZ deeper in the glycerol region, which in combination with sphingolipids may be involved in the lipid raft formation, leading to specific sorting and protein trafficking at the synaptic vesicle level, differentiating CLOZ from CPZ.

Overall, the demonstrated differences between CLOZ and CPZ in model membranes (Table 4) may contribute to the understanding of their distinct clinical and safety profiles, involving their influence on the GPCR lipid environment and on the lipid membrane properties of synaptic vesicles. For example, AP-induced changes of membrane fluidity and lipid raft composition may affect the distribution and function of dopamine receptors and synaptic vesicles, particularly their fusion properties, with differences due not only to their molecular structure but also to their accumulation overtime and protonation state. The biophysical membrane change differences observed between both APs may be associated with CLOZ treatment advantages when compared to CPZ.

**Table 4.** Comparison of membrane partitioning and impact in membrane properties of CPZ and CLOZ.

| Property/approach | CPZ | CLOZ |
| --- | --- | --- |
| Lipophilicity (logP) | Stronger (5.41) | Weaker (3.23) |
| Membrane partitioning (ITC) | Stronger | Weaker |
| Thermodynamics of partition (ITC) | -Enthalpy driven and higher for negatively charged membranes;<br>-Entropy driven for all systems except system 3 | -Enthalpy driven and higher (but less than CPZ) for negatively charged membranes;<br>-Entropy driven for all lipid systems |
| Binding reversibility (PWR) | No | Yes |
| Effect on zeta potential | Increase | Increase |
| Localization (AA-MD simulations) | Phosphate/glycerol level | Phosphate/glycerol level |
| Effect on headgroup region (Laurdan) | Ordering (weaker); strongest in system 1 and weakest in system 3 | Ordering (stronger); strongest in system 1 and weakest in system 3 |
| Effect on disordered membranes (AA-MD simulations, NMR) | Ordering | Ordering |
| Effect on ordered membranes (DSC, NMR) | Disordering (stronger) | Disordering (weaker) |
| Effect on membrane thickness (AFM) | Reduction | No effect |
| Membrane defects/pores (AFM, dye leakage) | No, expect at high ratios in system 3 only | No |
| Effect on vesicle size (DLS) | Reduction | Reduction |

The present study opens new avenues in the understanding of the non-receptor related actions of APs and contributes to the understanding of the recently proposed approaches of schizophrenia splitting the schizophrenia spectrum into different endophenotypic biotypes, leading to a more pathophysiological based prescription.^74^

## Supporting Information

The contents present in this section include: an experimental section with detailed information on Laurdan spectroscopy, particle size, zeta potential, isothermal titration calorimetry, sulforhodamine leakage, plasmon-waveguide resonance, ssNMR and simulation methods; 8 supplementary figures and 3 tables. “This material is available free of charge via the Internet at http://pubs.acs.org.”

## ACKNOWLEDGMENTS

Michael Molinari and the AFM platform in the CBMN are thanked for providing access to instrumentation. The NMR platform is thanked for instrument time. The ANR grant polyFADO financially support part of the studies. JB and JBK were funded in part by the National Science Foundation (NSF, US) with grant (CHE-2003912). The University of Maryland super-computing resources (http://hpcc.umd.edu) Zaratan were utilized for conducting the AA-MD simulations. Moreover, AAMD simulations were also carried out at the Advanced Research Computing at Hopkins (ARCH) core facility (rockfish.jhu.edu), which is supported by the NSF grant number OAC1920103 and time provided with allocation MCB100139 from the Advanced Cyberinfrastructure Coordination Ecosystem: Services & Support (ACCESS) program. GS thanks the National Science Fund of Bulgaria through Grant KP-06-H58/2021; the Bulgarian Ministry of Education and Science for the support provided via Scientific Infrastructure on Cell Technologies in Biomedicine D01-361/2023 and the National Center for Biomedical Photonics D01-352/2023, part of the Bulgarian National Roadmap for Scientific Infrastructures 2020–2027. MB acknowledges funding from the Portuguese Foundation for Science and Technology (FCT) to CIQUP, Faculty of Science, University of Porto (UIDB/00081/2020: DOI 10.54499/UIDP/00081/2020), and IMS-Institute of Molecular Sciences (LA/P/0056/2020). PN wishes to express his gratitude to the Fondation de France for the grant awarded to CFM (letter of commitment No. 00101826 in 2019).

## ABBREVIATIONS

(ΔH): Enthalpy change
(ΔS): Entropy change
(ΔG): Free Gibbs energy change
(AA-MD): All-Atom Molecular Dynamics
(AFM): Atomic Force Microscopy
(APs): Antipsychotics
(Chol): Cholesterol
(CLOZ): Clozapine
(CPZ): Chlorpromazine
(D2R): Dopamine D2 receptor
(DLS): Diffraction Light Scattering
(DSC): Differential Scanning Calorimetry
(GP): General Polarization
(GPCRs): G-Protein Coupled Receptors
(ITC): Isothermal Titration Calorimetry
Kp): Partition constant
(LDCVs): Large Dense Core Vesicles
(LUVs): Large Unilamellar Vesicles
(MLVs): Multilamellar Vesicles
(PWR): Plasmon-Waveguide Resonance
(SLB): Supported Lipid Bilayer
(ssNMR): SolidState NMR
(SM): Sphingomyelin
(SVs): Synaptic Vesicles

## Table of Contents

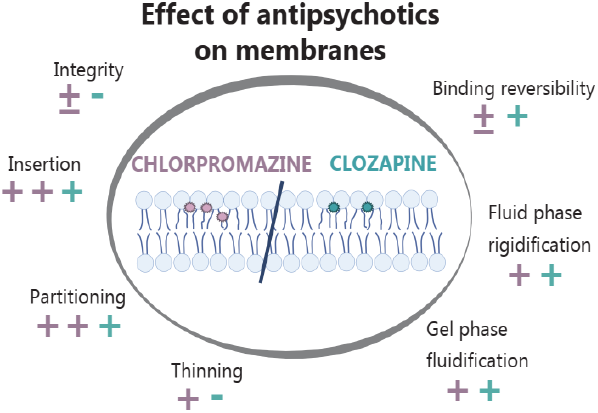

## Supplementary Information

## Experimental section

### Preparation of Large Unilamellar Vesicles (LUVs)

#### a. For Laurdan spectroscopy, particle size and zeta potential

The lipids were dissolved and mixed in chloroform to yield a concentration of 1 mM. Appropriate volumes of stock solutions of APs (1 mg/mL) were added to the lipids in the initial organic solutions. The solvent was removed under a stream of oxygen-free dry nitrogen and the residues were left in a vacuum desiccator overnight at room T. Then the obtained lipid films were hydrated with DPBS (1x), 150 mM NaCl, pH 7.2. The hydrated lipids were subjected to three cycles of vortex/sonication at room temperature in case of system 2 and 3. As for system 1, samples were heated at 60 °C for 10 min, vortexed for 1 min, then left in a sonication bath for 1 min, and finally cooled in ice for 5 min. This heating/cooling procedure was repeated three times to ensure sample homogenization. MLVs produced at this stage were extruded with a LiposoFast small-volume extruder (Avestin, Ottawa, Canada) equipped with Nuclepore™ polycarbonate track-etched membrane filters (Cytiva’s Whatman™) as follows: 11 extrusions through 800 nm, followed by 21 extrusions through 100 nm filters to obtain LUVs. For the extrusion steps for system 1, the aqueous MLV suspensions were maintained at 60 °C. Each sample was measured in final lipid concentration of 200 µM. LUV samples were studied the same day after incubation for 1 h at 37 °C. To check the concentrations of LUVs after the extrusion the total amount of phosphorous was assessed in a LUV sample following a modified procedure of Fiske & Subbarow from Avanti Polar Lipids.^1^ The predicted concentrations of LUVs were found to be accurate within 5 %.

#### b. For isothermal titration calorimetry measurements

The lipid films were hydrated with HEPES 25 mM, KCl 150 mM, pH 7.4. The MLVs formed were then subjected to five freeze-thaw cycles (liquid nitrogen - water bath 60 °C). The dispersions were extruded 11-times through 200 nm Nucleopore™ Track-Etched membrane using Avanti polar lipid extruder at room T. After extrusion, Rouser method for determination of total phosphorous was applied to obtain the LUV concentration before the experiment.^2,3^

#### c. For sulforhodamine B (SRB) leakage

The lipid concentration for sulforhodamine B (SRB) leakage experiments was 2 mg/mL. After lipid film preparation, hydration was performed by adding buffer HEPES 25 mM, KCl 50 mM, pH 7.4, containing 50 mM of fluorescent dye SRB. The freeze-thaw cycles and extrusion were performed as described for ITC lipid procedure, and then the vesicles were placed onto Sephadex G-75 size exclusion column equilibrated with buffer in order to separate SRB encapsulated vesicles from the free dye. The elution buffer used was HEPES 25 mM, KCl 150 mM, pH 7.4. The phospholipid determination of the vesicles allowed to measure the concentration of LUVs present in the sample liberated from free dye, following the Rouser method.^2,3^ The measurements were performed on Tecan at 25 °C (λ_ex_ and λ_em_ used were 555 and 577 nm, respectively) on 96 black flat bottom well-plates. The vesicles were diluted to 140 µM and 50 µL were added to each well. Then, 50 µL of APs (CPZ or CLOZ) at appropriate concentrations to reach the different lipid:AP ratios (100:1, 50:1, 25:1 and 10:1) were added in separate wells. The buffer used for preparation of AP solutions was the same as for dye-encapsulated liposomes (column elution buffer). Fluorescence intensity measurements were taken following 1 h liposome:AP incubation. Maximal leakage was induced by adding 100 µL of 1 % Triton X-100, which completely dissolved the liposomes. At least three measurements were made with three separate liposome preparations. The following equation was used for calculation of leakage:

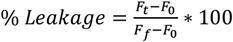

The F_0_ represents the fluorescence intensity of vesicles without AP at time 0, the F_t_ is the fluorescence at time t (1 h), and the F_f_ is the final fluorescence after Triton X-100 was added. This technique allows the measurement of leakage percentage of SRB fluorophore upon a potential effect of AP on lipid membrane integrity (formation of pores).

### Plasmon-waveguide resonance

Like classical surface plasmon resonance (SPR), plasmon-waveguide resonance (PWR) uses a polarized continuous wave laser to excite the surface plasmons in a resonator made of a thin metallic film deposited on a prism front face. The major difference lies in the additional presence of a thick silica layer - acting as a waveguide – coating the silver layer that allows to access not only the resonance in *p*-polarized light (parallel to the incident light, *i*.*e*. perpendicular to the sensor) but also in *s*-polarized light (perpendicular to the incident light, *i*.*e*. parallel to the sensor), with much higher sensitivity.^4^ The molecules attached at the outer surface of silica interact with the evanescent field and subsequently change the resonance characteristics, which also depend on the incident angle of the laser beam. The changes in the amplitude, position, and width of the reflected light intensity as a function of the incident angle translate into optical and mechanical parameter variations illustrating, *e*.*g*., a change in mass, thickness or refractive index of the immobilized and/or interacting materials. Such variations can be followed in real-time and as a function of concentration to follow the kinetics and thermodynamics properties of molecular interactions.

PWR spectra were obtained with a home-made instrument equipped with a He-Ne laser at 632 nm, allowing the acquisition of both *p*- and *s*-polarization of the light, a prism sensor coated with a 460 nmsilica layer on top of a 50 nm - silver layer and a rotating table allowing incremental changes of 1 mdeg for the incident light. The reflected light as a function of the incident light is detected by a photodiode (Hamamatsu) and registered on a photodiode amplifier (Thorlabs). The setup is controlled *via* a homemade LabView program, allowing to monitor, in real time, not only the raw spectra but also critical parameters such as the resonance minimum position for *p-* and *s-*pol.^5^

The SUV lipid suspension (3 mg/mL in HEPES 25 mM, KCl 150 mM, pH 7.5) is introduced in a Teflon block (V = 250 µL) in close contact with the sensor, so that the lipid bilayer can form at the prism interface. Its formation is followed in real time by the changes in the resonance angles in *p-* and *s-*polarizations. Excess of lipid solution is then removed by washing the bilayer with buffer. Once the bilayer is formed and stable, molecules of interest (either CPZ or CLOZ) are injected in the Teflon block, in an incremental manner (from sequentially diluted solutions 0.001 µM to 100 µM in the same buffer as SUVs), to follow in real time its effect on the bilayer. Before each injection, a time delay of 10 min is respected to reach a saturation effect at a defined concentration.

### Solid-state NMR

One lipid composition was used: pure 1,2-dipalmitoyl-(^2^H_62_) oleoyl-*sn*-glycero-3-phosphocholine (^2^H_62_-DPPC) as a simplistic model. Sample was prepared by premixing lipid with AP in chloroform (ratio lipid:AP 9:1). After solvent evaporation, the film was dispersed in 1 mL milliQ filtered water and freeze-dried overnight to remove all traces of the solvent. The resulting powder sample was suspended in 100 µL of deuterium-depleted water to obtain 90 % hydration. Three freeze-thaw cycles were repeated (the samples were well vortexed between each cycle for better sample homogeneity): the samples were frozen in liquid nitrogen before being heated at 40 °C for 10 min, and the resulting milky dispersion was transferred into a 4 mm diameter Zirconium NMR rotor.

Solid-state ^2^H NMR measurements were performed on a Bruker Avance III 500 MHz SB spectrometer (Bruker Biospin, France) equipped with a dual 1H/X 4-mm magic ancle spinning (MAS) probe. ^2^H-NMR spectra were acquired at 76.77 MHz by means of a quadrupolar echo pulse sequence with a spectral width of 500 kHz, a π/2 pulse width of 3.5 μs, a 50 μs interpulse delay, 4k scans, a time domain of 2K points and a recycle delay of 2 s.^6^ A Lorentzian noise filtering of 300 Hz was applied prior to Fourier transformation from the top of the echo signal.

Samples were allowed to equilibrate during at least 20 min at a given temperature before the NMR signal was acquired.

The processing of the ^2^H NMR experimental spectra gives access to several parameters describing the dynamics of the membrane: quadrupolar splitting Δν_Q_, order parameter S_CD_ and the first moment M_1_. Alternatively, spectra can be calculated. Processing and simulations of the NMR data were conducted as previously described by Beck et al. and Grélard et al.^7,8^

### Simulation methods

#### Parameterization of CPZ and CLOZ

The ligands CPZ and CLOZ, protonated and uncharged, were parametrized with the CGENFF module of CHARMM-GUI.^9,10^ To validate the accuracy of the parameterization, the logP of lipids for partitioning between water and octanol was estimated. To obtain the octanol/water logP of the ligands, three replicate simulations of the ligands were done in pure water and octanol (73.4 %)-water (26.6 %) boxes using GROMACS. The ligand configurations for the replicates were obtained from short production runs of the ligand in a 30 Å water box. The ligand was placed in the center of a box with 818 TIP3 water molecules, energy minimized using steepest descent and equilibrated in NVT for 125 k steps with 1 fs timestep and with a harmonic restraint of force constant 400 kJ mol^-1^ nm^-2^ applied on the ligand.^11^ This was followed by a production run of 500 k steps and 2 fs timestep in NPT without restraints. The Nose-Hoover thermostat was used for maintaining the temperature and the Parinello-Rahman barostat with isotropic coupling was used for the ligand and solvent. Configurations for the three replicates were extracted from the production run to set up the free energy perturbation (FEP) systems. FEP was done for both ligands in CHARMM to get the free energy of partitioning (water/octanol) based on the current forcefield parameters.^12^ Each ligand-solvent system had three sets of simulations for the three non-bonded interactions: Lennard-Jones attractive, repulsive interactions and electrostatic charged interactions. They were energy minimized for a total of 400 steps with the Steepest Descent (SD) and Adopted Basis Newton-Raphson (ABNR) methods with a harmonic restraint of 100×10^6^ kCal/mol applied to non-hydrogen atoms. The systems were then equilibrated for 50 k steps with a time-step of 2 fs at temperature of 298.15 K. The production runs were integrated using the new velocity Verlet (vv2) method for 100 k steps with 2 fs timestep. The switched scheme with 1-1.2 nm cutoff was used for the van der Waals interactions, Particle Mesh Ewald summation with spline order 6 and kappa 0.34 was used for electrostatic ones. The ligand was restrained within a sphere of radius 1 nm centered at the origin. The Geometric module from CHARMM’s Miscellaneous Mean Field Potential (MMFP) was used to apply a harmonic potential with force constant 1 kCal/mol on the ligand, whenever it extended beyond the spherical region. For the free energy calculations, CHARMM’s Perturb module was used to scale the non-bonded interactions in each set of simulation. The interactions were independently scaled by factor 1-λ, where λ was incremented from 0 to 1 with step-size 0.1. Thus, they were scaled (1-λ) from 1 (fully interacting) down to zero (noninteracting). CHARMM’s implementations of the Weighted Histogram Analysis Method (WHAM) and Thermodynamic Integration (TI) were used to combine and extract the free energy of the perturbation. The two methods gave almost identical results and the greater value of the two were selected. Since the calculated octanol/water partition free energy is close to the experimental value, we used the CGENFF parameterization without any further changes.

### System setup and simulation protocol

#### Membrane-only

The all-atom simulations were done with namd 2.14 and GROMACS 2021.4 using the CHARMM36 forcefield.^13–17^ First, the bilayer-only systems were constructed for three membrane compositions. CHARMM-GUI’s *Membrane Builder* was used to get the three different bilayers solvated in water and with neutralizing NaCl ions.^18,19^ Systems 1 and 3 were simulated using namd, while system 2 was simulated using GROMACS and was energy minimized for 5 k steps using steepest descent, followed by two NVT (constant Number of particles, Volume, and Temperature) equilibrations of 125 k, 125 k steps with 1 fs timestep and with restraints on the lipid backbone and sidechains. Four NPT (constant Number of particles, Pressure, and Temperature) equilibration steps were done of 125 k, 250 k, 250 k and 250 k steps with 1 fs, 2 fs, 2 fs and 2 fs timestep, respectively. All the equilibration steps used the Berendsen thermostat to maintain the temperature of the system at 296.15 K using a τ_t_ of 1 and the Berendsen barostat with semi-isotropic coupling with a compressibility of 4.5 x 10^-5^ and τ_p_ of 5 to maintain 1 bar pressure. The NPT production runs used the Nosé-hoover thermostat and Parinello-Rahman barostat with semi-isotropic coupling and the LINCS algorithm was used to restrain hydrogen bonds. The systems 1 and 3 were first energy minimized and equilibrated in NVT and then in NPT in namd with the same standard protocol as above. They were energy minimized for a maximum of 10 k steps with a timestep of 1 fs. Langevin dynamics at temperature 296.15 K with restraints on the lipids was used. A constant ratio along the x-y box vector was maintained for the membrane volume. This was followed by another 125 k steps of equilibration in NVT ensemble. Equilibration in NPT ensemble was done the same as for system 2. Finally, the production runs were done in the NPT with the lipid restraints removed. The pressure was maintained at 1 atm using a Langevin piston with an oscillation period and decay of 50 fs, 25 fs, respectively. All simulations used the van der Waals force switch scheme with 1-1.2 nm cutoffs and Particle Mesh Ewald (PME) summation with a radius of 1.2 nm for the electrostatics and the Verlet cutoff scheme was used for the neighbor’s list with a cutoff radius of 1.2 nm. The membrane composition and total simulation times are listed in Table S1.

#### Membrane-AP

The pre-equilibrated membranes and the APs (CPZ or CLOZ) were used to build the systems of interest with the help of CHARMM-GUI’s *Muti-component assembler* module.^20^ Three systems were built, system 1-3. Each membrane system was symmetric and simulated in a low (100:9) and high (100:20) lipid:ligand ratio per leaflet. The membrane composition and total simulation times are listed in Tables S2 and S3.

The membrane-ligand system was energy minimized and equilibrated using the standard CHARMM-GUI protocol and methods as mentioned earlier. Energy minimization was done using steepest descent with the default restraints on the lipid backbone and sidechains, along with an additional restraint of force constant 5000 kJ mol^-1^ nm^-2^ on the ligands to hold them in position. Following this, two equilibration steps in the NVT ensemble were done with the same schemes as mentioned before. The next three equilibration steps were done in NPT ensemble with the Berendsen barostat with semi-isotropic coupling, because of the membrane. The force constant of the restraints on the lipids were gradually reduced in successive equilibration steps based on the standard CHARMM-GUI protocol, while the restraint on the ligands were kept constant. The production runs were done in the NPT ensemble with a timestep of 2 fs with the same scheme as mentioned before.

### All-atom simulation analysis

The all-atom ligand membrane simulations were analyzed for hydrogen order parameter (S_CH_), electron density, area per lipid and z-coordinate of the ligands. The S_CH_ value of the lipid chains indicate the chain flexibility. This can change due to the presence of certain membrane-bound molecules or due to a change in lipid packing. The S_CH_ is defined as follows:

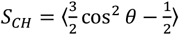

where θ represents the angle between the C-H bond vector and the membrane normal. Thus, when the C-H bond is parallel to the membrane normal, the S_CH_ is the highest (1) and there is tight packing between the lipids making them “gel-like”. However, when the C-H bond is perpendicular to the membrane normal, the S_CH_ is the lowest (0.5) and the lipid packing is not compact making them “fluid”. Another parameter is the electron density of the lipids. Using the electron density, we can measure the membrane thickness and the position of the lipid functional groups along the membrane normal. It is also helpful to look at the ligand distribution along the membrane normal to get an estimate of the ligand’s preference for the membrane-surface, membrane-interior or the bulk water. The Cl atom of both CPZ and CLOZ were used as a proxy for the aromatic moieties of the ligands. The area per lipid versus time is a metric that can be used to determine whether the membrane system has equilibrated. For the small systems simulated, the area per lipid is just the product of the *X* and *Y* cell dimension divided by the number of lipids per leaflet. It also gives us a clue about the lipid packing since lower area per lipid indicates that the lipids are well-packed and this could make it harder for any molecule to penetrate the membrane.

## Supplementary Figures

**Figure S1.**
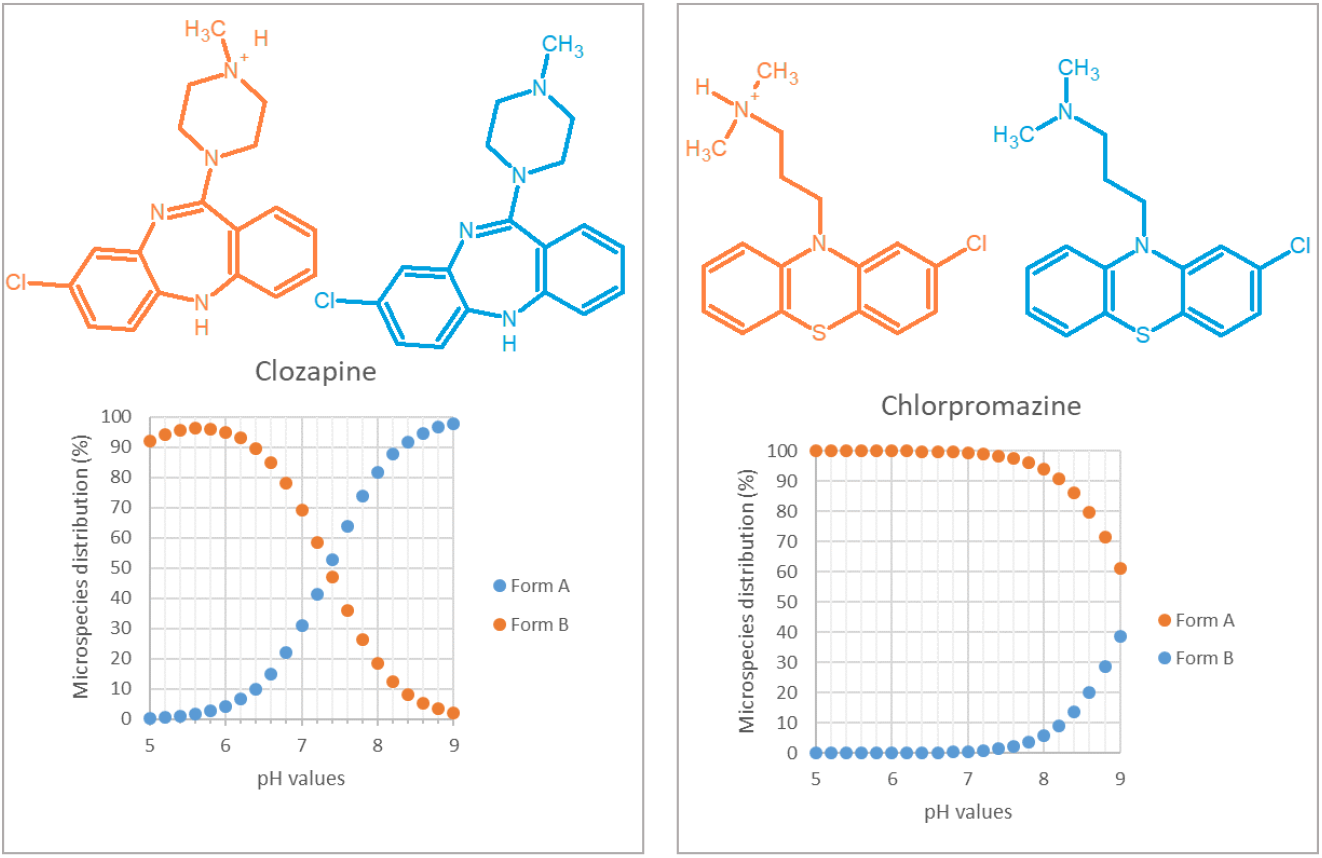
Marvin Sketch simulations of the distribution of non-protonated (form A, blue) and a protonated (form B, orange) species in terms of pH is represented for CLOZ (left) and CPZ (right) (Chemaxon). Experimental pKa values are 7,5 and 9,3 for CLOZ and CPZ, respectively.^21,22^

**Figure S2.**
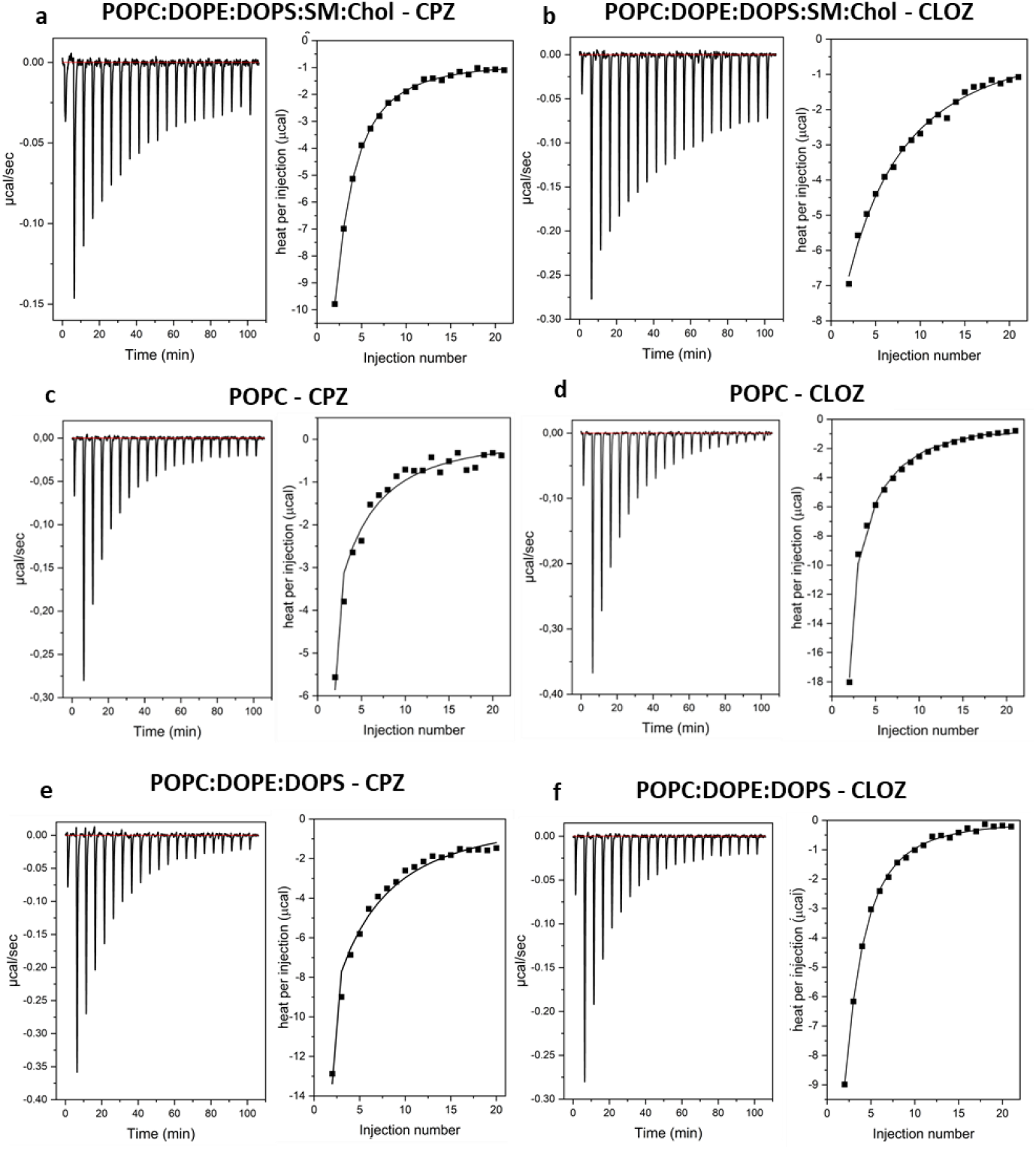
ITC thermograms (left) with the corresponding titration curves (right) – pH 7.4, 25 °C. System 1 is represented with CPZ (a) and CLOZ (b), as well as system 2: with CPZ (c) and CLOZ (d) and system 3: with CPZ (e) and CLOZ (f).

**Figure S3.**
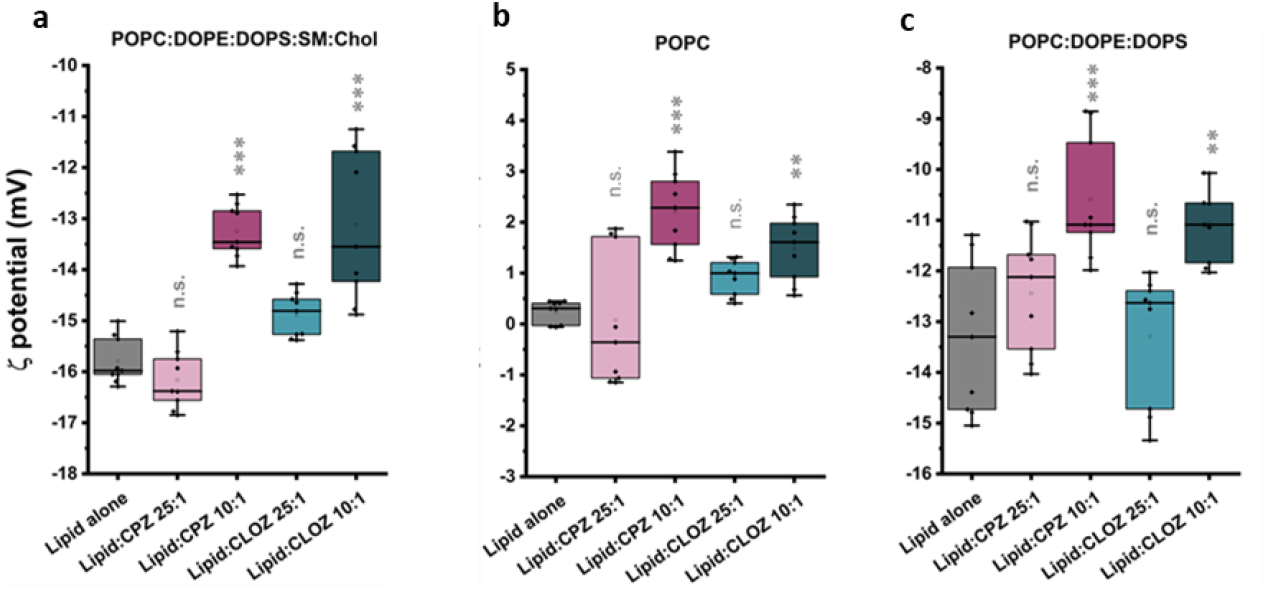
ξ potential of control LUVs without AP (grey) and AP-containing ones (CPZ – pink, CLOZ - cyan) in lipid:AP ratios 25:1 and 10:1. Lipid systems 1 (a), 2 (b) and 3 (c) have been used for the measurements. One way ANOVA test has been used for comparing each dataset with the control group (lipid alone, the results are shown above each box).

**Figure S4.**
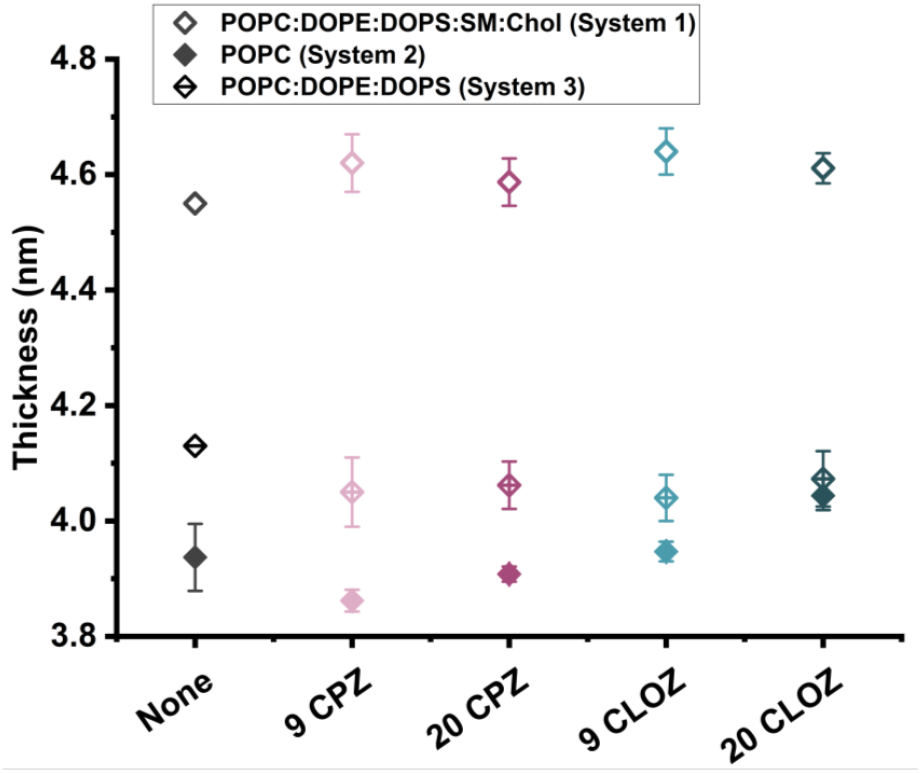
Bilayer thickness measured from P-P atom distance of membrane leaflets for system 1-3 with and without AP presence (either 9 or 20 molecules of each AP) with AA-MD simulations. Membrane alone is represented in grey, CPZ in pink and CLOZ in cyan.

**Figure S5.**
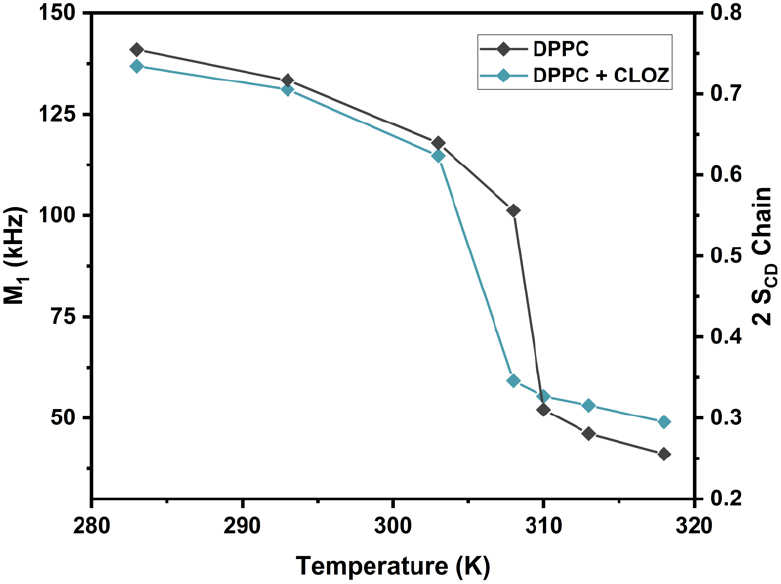
First spectral moments (M_1_), and corresponding whole chain ordering (2S_CD_chain, double y-axis) as a function of temperature for ^2^H_62_-DPPC in absence (black) and presence (blue) of CLOZ at lipid:AP molar ratio 10:1.

**Figure S6.**
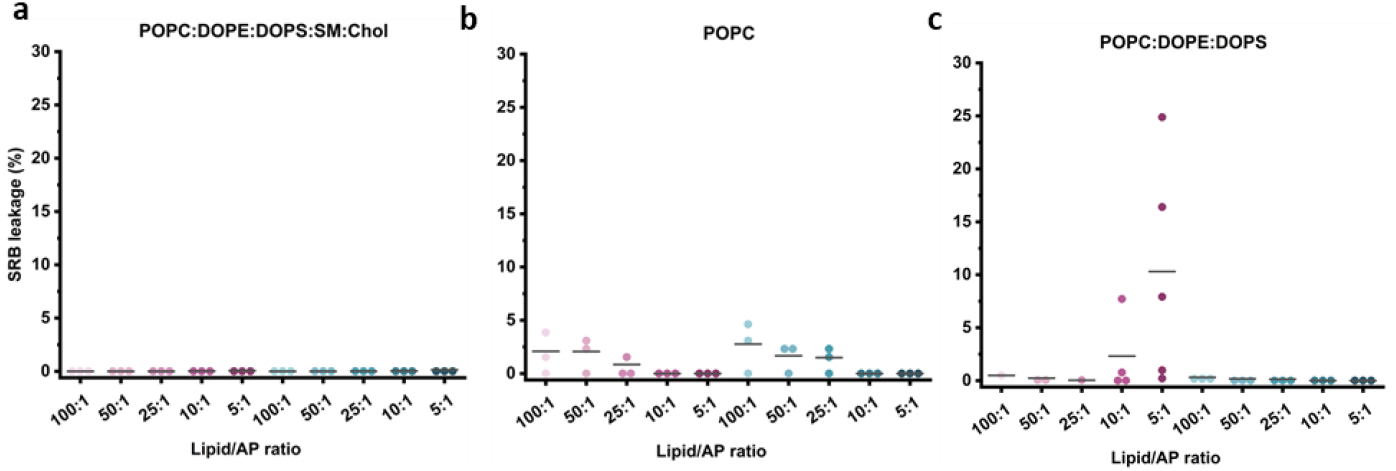
Dye leakage (%) after 1 h incubation of liposomes encapsulating sulforhodamine in absence and presence of the two AP at lipid:AP molar ratios of 100:1, 50:1, 25:1, 10:1 and 5:1. CPZ is shown in pink and CLOZ in cyan. Three different lipid systems have been investigated: system 1 (a), system 2 (b) and system 3 (c).

**Figure S7.**
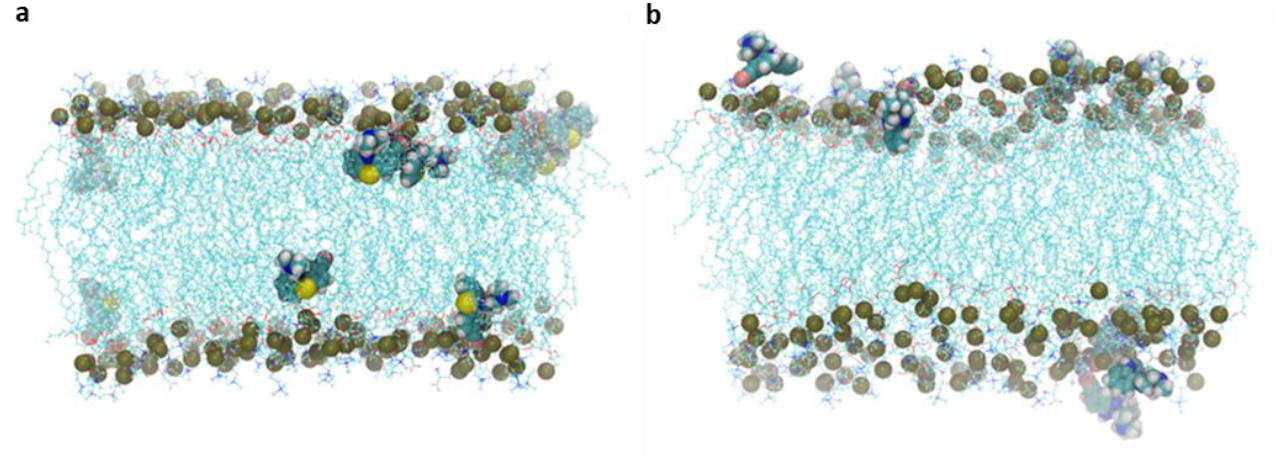
Representative snapshot of system 3 with 9 CPZ (a) and 9 CLOZ (b) interacting with and/or embedded in the membrane, obtained with AA-MD simulations.

## Supplementary Tables

**Table S1.**
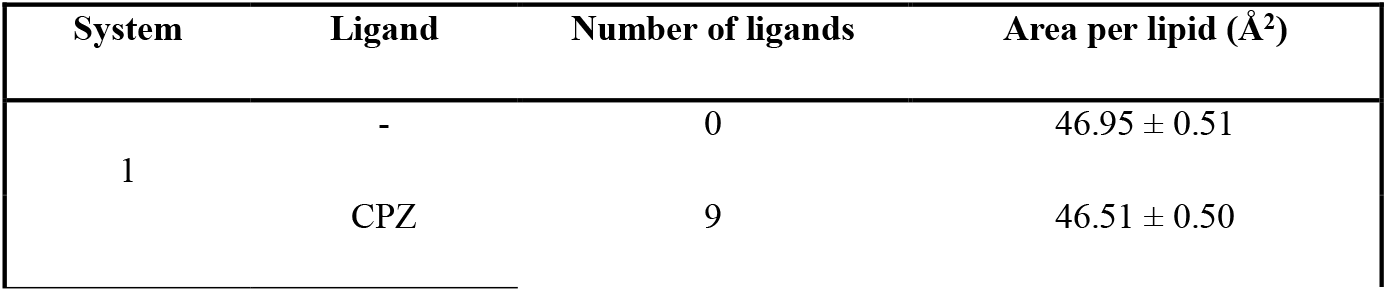
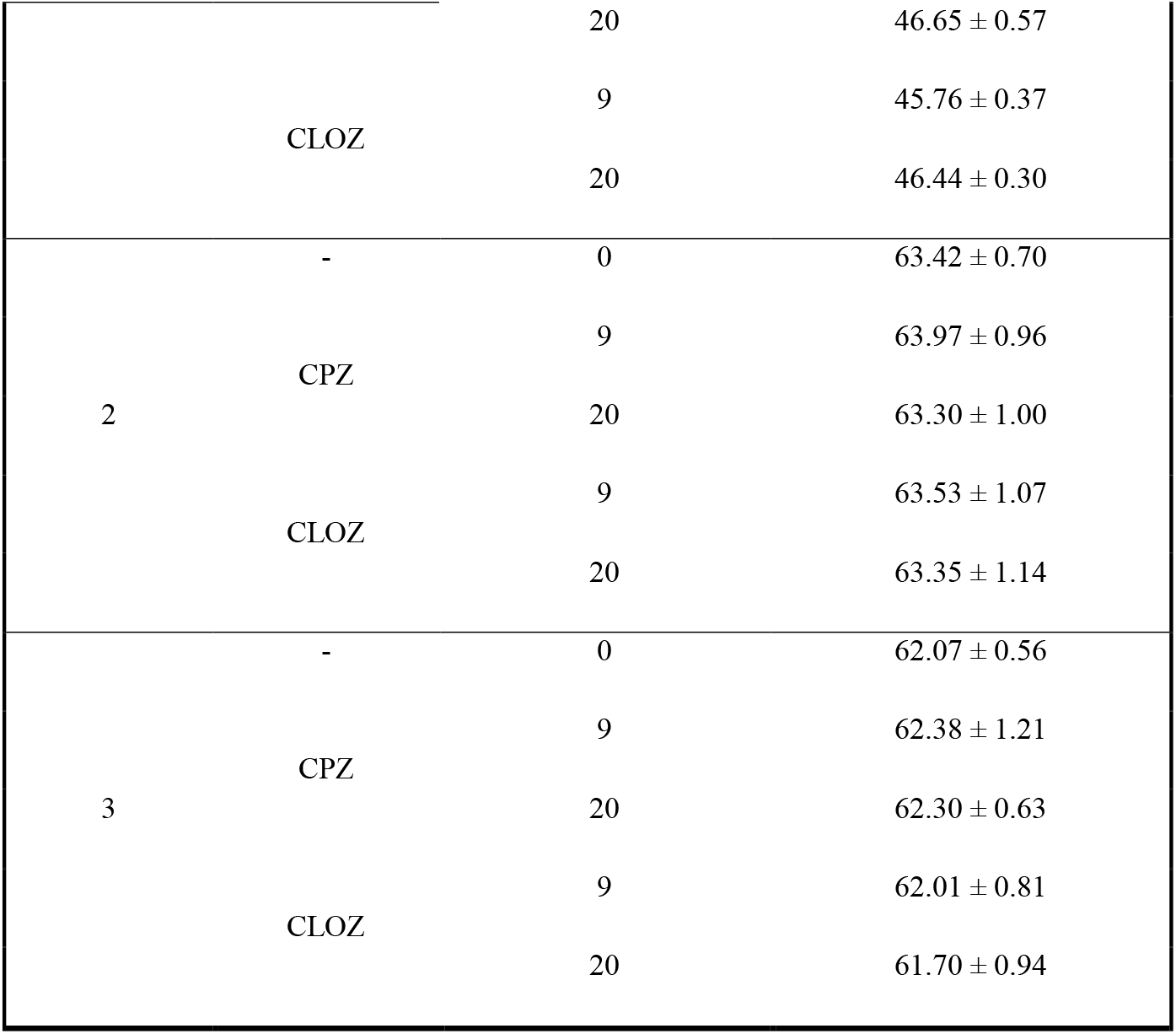
Area per lipid for the three systems in absence/presence of either 9 or 20 CPZ/CLOZ obtained by AA-MD simulations.

**Table S2.** Membrane only system composition and simulation time (AA-MD simulations).

| System | #POPC | #DOPS | #DOPE | #CHOL | #SM | #TIP3 | Simulation<br>time (ns) |
| --- | --- | --- | --- | --- | --- | --- | --- |
| 1 | 200 | 0 | 0 | 0 | 0 | 9040 | 400 |
| 2 | 50 | 20 | 50 | 60 | 20 | 7680 | 267 |
| 3 | 130 | 20 | 50 | 0 | 0 | 8931 | 150 |

**Table S3.**
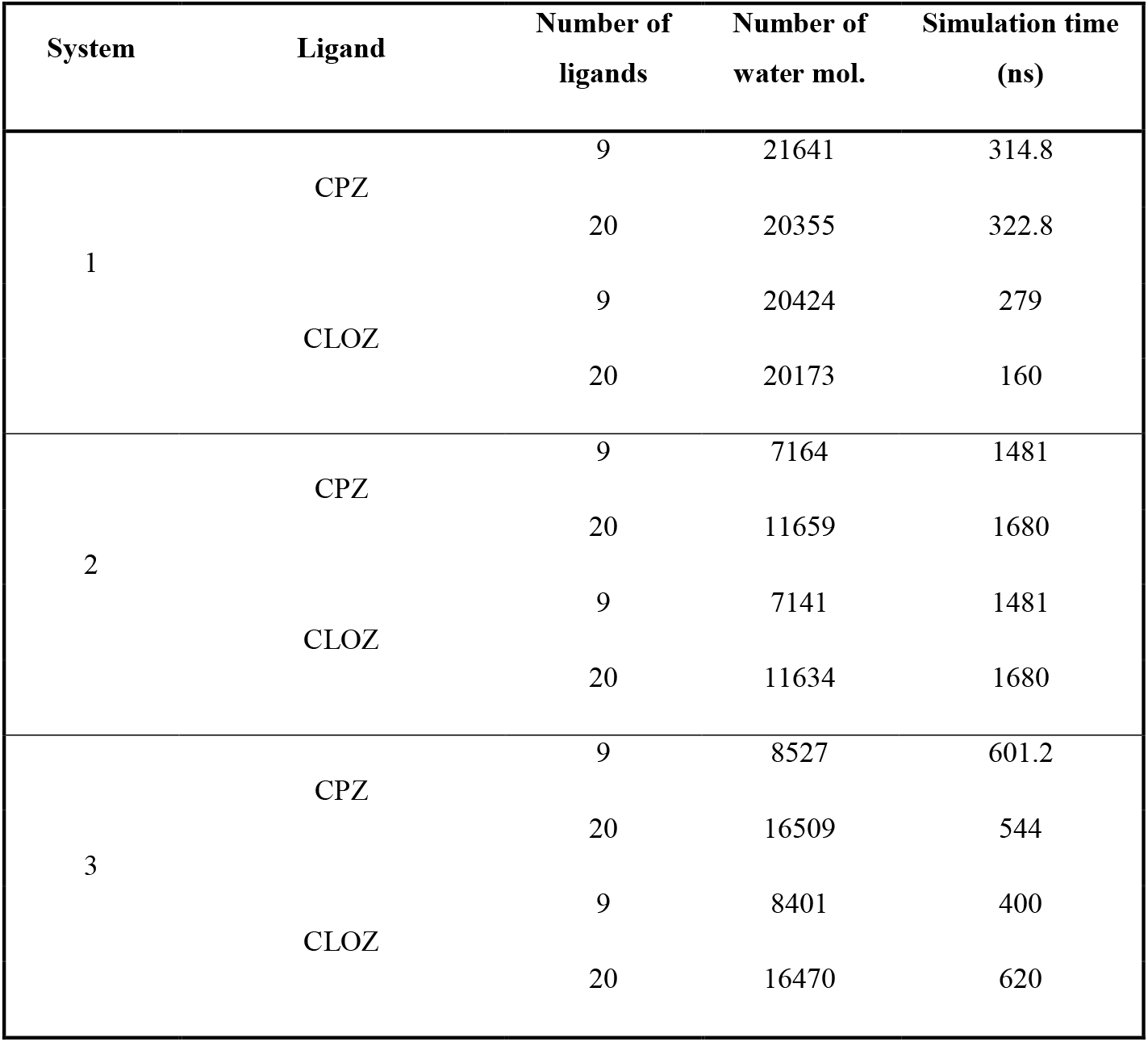
Membrane system composition and simulation times (AA-MD simulations).

**Table S4.** Thermodynamic parameters obtained by ITC for the interaction and partition of CLOZ and CPZ with liposomes with systems 1, 2 and 3. Three separate measurement done for each system.

| System | AP | $K_P (10^4)$ | $\Delta H (kJ mol^{-1})$ | $\Delta G (kJ mol^{-1})$ | $\Delta S (J K^{-1} mol^{-1})$ |
| --- | --- | --- | --- | --- | --- |
| POPC:DOPE:DOPS:SM:Chol | CPZ | $1.4 \pm 0.1$ | $-18.2 \pm 2.7$ | $-23.6 \pm 0.2$ | $19.2 \pm 9.2$ |
| | CLOZ | $0.08 \pm 0.03$ | $-5.5 \pm 2.2$ | $-16.6 \pm 0.8$ | $37.3 \pm 4.7$ |
| POPC | CPZ | $1.3 \pm 0.2$ | $-12.9 \pm 1.6$ | $-23.5 \pm 0.4$ | $35.5 \pm 6.5$ |
| | CLOZ | $0.32 \pm 0.12$ | $-6.0 \pm 2.6$ | $-19.9 \pm 1.1$ | $46.7 \pm 5.2$ |
| POPC:DOPE:DOPS | CPZ | $1.3 \pm 0.0$ | $-29.4 \pm 4.6$ | $-23.5 \pm 0.0$ | $-19.7 \pm 15.7$ |
| | CLOZ | $0.38 \pm 0.00$ | $-10.0 \pm 1.1$ | $-20.4 \pm 0.0$ | $34.9 \pm 3.6$ |

**Table S5.** Z-average and polydispersity index (PDI) of LUVs with and without AP, analyzed by DLS. LUVs have been prepared using lipid systems 1, 2, and 3 (concentration 0.2 mM), with lipid:AP ratios of 25:1 and 10:1. Measurements have been conducted at 37 °C, with a minimum of three replicates performed on each of three independently prepared samples. Data are presented as means ± SD (n = 9).

| System | AP | lipid:AP ratio | Z-average, nm | PDI |
| --- | --- | --- | --- | --- |
| 1 | - | 0 | 135.4 ± 1.9 | 0.037 ± 0.015 |
|  | CPZ | 25/1 | 133.4 ± 1.5 | 0.047 ± 0.033 |
|  |  | 10/1 | 132.9 ± 1.0 | 0.071 ± 0.050 |
|  | CLOZ | 25/1 | 130.6 ± 1.3 | 0.053 ± 0.035 |
|  |  | 10/1 | 128.6 ± 1.3 | 0.068 ± 0.040 |
| 2 | - | 0 | 138.9 ± 0.7 | 0.075 ± 0.060 |
|  | CPZ | 25/1 | 136.2 ± 0.9 | 0.082 ± 0.016 |
|  |  | 10/1 | 139.6 ± 1.3 | 0.078 ± 0.034 |
|  | CLOZ | 25/1 | 134.8 ± 1.3 | 0.090 ± 0.026 |
|  |  | 10/1 | 133.3 ± 1.8 | 0.069 ± 0.030 |
| 3 | - | 0 | 130.8 ± 1.5 | 0.080 ± 0.044 |
|  | CPZ | 25/1 | 126.1 ± 1.3 | 0.080 ± 0.034 |
|  |  | 10/1 | 130.8 ± 0.9 | 0.064 ± 0.037 |
|  | CLOZ | 25/1 | 130.7 ± 1.8 | 0.044 ± 0.019 |
|  |  | 10/1 | 129.1 ± 1.4 | 0.085 ± 0.021 |

